# Nicotinate degradation in a microbial eukaryote: a novel, complete pathway extant in *Aspergillus nidulans*

**DOI:** 10.1101/2021.08.17.456622

**Authors:** Eszter Bokor, Judit Ámon, Mónika Varga, András Szekeres, Zsófia Hegedűs, Tamás Jakusch, Michel Flipphi, Csaba Vágvölgyi, Attila Gácser, Claudio Scazzocchio, Zsuzsanna Hamari

## Abstract

Several strikingly different aerobic and anaerobic pathways of nicotinate utilization had been described in bacteria. No similar work is extant in any eukaryote. Here we elucidate a complete eukaryotic nicotinate utilization pathway, by constructing single or multiple gene deleted strains and identifying metabolic intermediates by ultra-high performance liquid chromatography – high-resolution mass spectrometry. Enzymes catalyzing each step and all intermediate metabolites were identified. We previously established that the cognate eleven genes organized in three clusters constitute a regulon, strictly dependent on HxnR, a pathway-specific transcription factor. The first step, hydroxylation of nicotinic acid to 6-hydroxynicotinic acid is analogous to that occurring in bacterial pathways and is catalyzed by an independently evolved molybdenum-containing hydroxylase. The following enzymatic steps have no prokaryotic equivalents: 6-hydroxynicotinic acid is converted to 2,3,6-trihydroxypyridine through 2,5-dihydroxypiridine and the trihydroxylated pyridine ring is then saturated to 5,6-dihydroxypiperidine-2-one followed by the oxidation of the C6 hydroxyl group resulting in 3-hydroxypiperidine-2,6-dione. The latter two heterocyclic compounds are newly identified cellular metabolites, while 5,6-dihydroxypiperidine-2-one is a completely new chemical compound. Ring opening between C and N results in α-hydroxyglutaramate, an unprecedented compound in prokaryotic nicotinate catabolic routes. The pathway extant in *A. nidulans*, and in many other ascomycetes, is different from any other previously analyzed in bacteria. Our earlier phylogenetic analysis of Hxn proteins together with the complete novel biochemical pathway we now describe further illustrates the convergent evolution of catabolic pathways between fungi and bacteria.

**Significance Statement:** This eukaryotic nicotinate catabolic pathway illustrates the convergent evolution of prokaryotic and microbial eukaryotic metabolism. It brings to light newly identified metabolites and step processing enzymes. The identification of hitherto undescribed metabolites - which could serve as precursor biosynthetic molecules - is potentially relevant to both pharmaceutical and agrochemical industries.

## Introduction

Nicotinic acid (niacin, vitamin B3), a precursor of NAD, can serve as a nitrogen and carbon source in bacteria. In prokaryotes nicotinic acid (NA) is first converted to 6-hydroxynicotinic acid (6-NA), a reaction catalyzed by MOCO (molybdenum cofactor)-containing nicotinate hydroxylase enzymes (reviewed in (1)), which evolved several times independently (2-4). Four quite different pathways have been described in detail in bacteria (5).

The only detailed study of nicotinate utilization in a eukaryotic microorganism was carried out by us in the ascomycete *Aspergillus nidulans*. A nicotinate hydroxylase was characterized, and mutants in a gene encoding this enzyme and a putative transcription factor necessary for its induction were described (6-10). The genes encoding nicotinate hydroxylase (HxnS) and the HxnR transcription factor map in a six-gene co-regulated cluster (including also *hxnZ,Y,P* and *T*, cluster hxn1/VI) (10). Recently, five additional *hxn* genes (*hxnX,W,V,N* and *M*) were identified as members of the HxnR-regulon. In *A. nidulans*, these map in two additional gene clusters (hxn2/VI and hxn3/I clusters) (11) (Fig. 1).

**Fig. 1.**
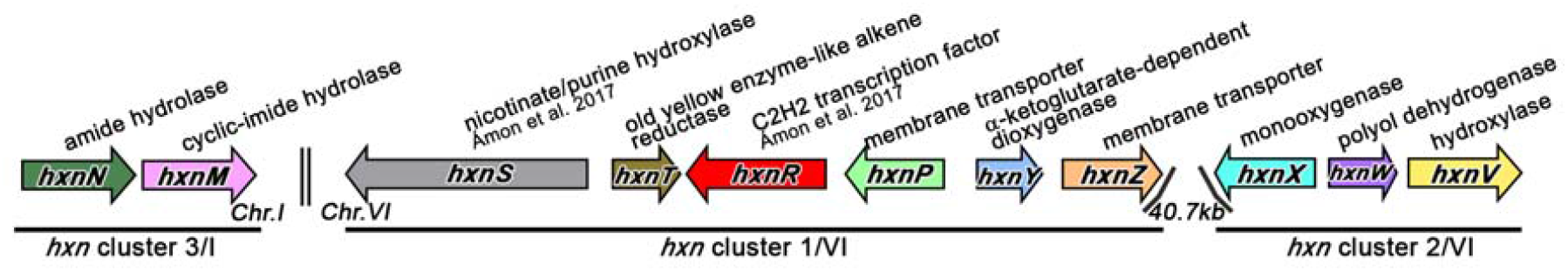
Organization of HxnR regulon in three gene clusters in *A. nidulans (11)*. Color arrows indicate specific *hxn* genes and relative gene orientation. Double vertical line symbolizes location of genes on different chromosomes. Above the coding genes, reported roles of gene products or roles deduced from domain functions are indicated.

All the *hxn* genes are induced by a hitherto non-determined derivative of nicotinic acid (further referred as physiological inducer) (10). Induction necessitates both the pathway-specific transcription factor HxnR and the GATA factor AreA, mediating nitrogen metabolite de-repression (10, 11). The *hxnR* gene is characterized by both loss of function (including deletions) and constitutive mutants (10).

In *Aspergillus terreus*, an RNASeq study determined that growth in the presence of salicylate results in induction of *hxnS* and *hxnX* orthologues through 3-hydroxyanthranilate-coupled quinolinate degradation (12). This suggests that in this organism, either a common inducer metabolite occurs in the nicotinate and salicylate degradation pathways, or that in the latter pathway a different metabolite can act as a positive effector of HxnR, too.

In this work, we establish the complete nicotinate degradation pathway in the ascomycete filamentous fungus *A. nidulans* by using reverse genetics and by ultra-high performance liquid chromatography – high-resolution mass spectrometry (UHPLC-HRMS) based analysis of pathway metabolites, followed by purification and NMR analysis of two compounds. This work illustrates the convergent evolution of metabolic pathways in phylogenetically very distant microorganisms.

## Results

Fig. 2 shows the pathway of NA utilization in *Aspergillus nidulans*. The rationale for this pathway is detailed below.

**Fig. 2:**
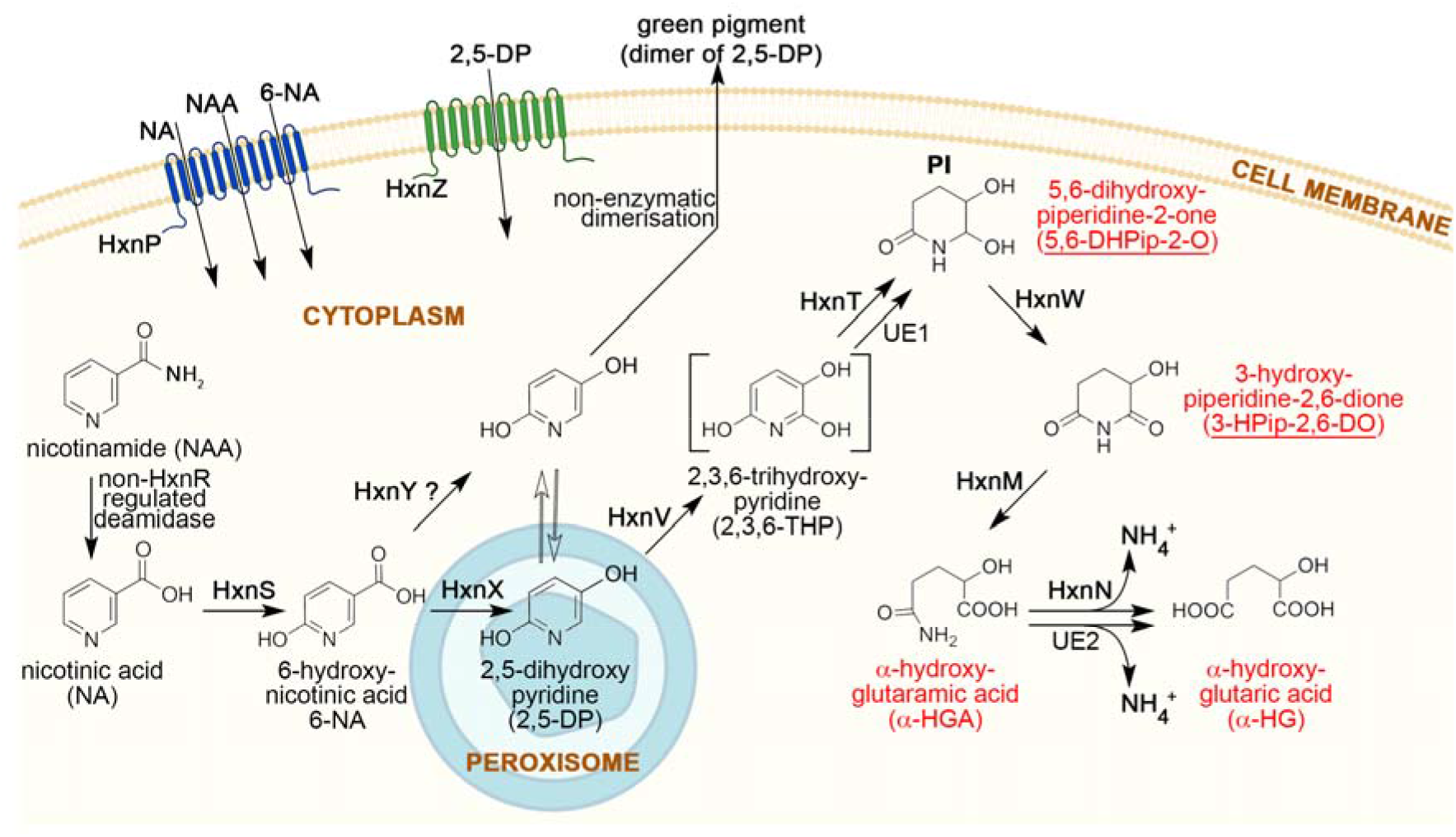
Nicotinate catabolic route in *A. nidulans*. HxnP and HxnZ are transporters (marked with blue and green transmembrane domains, respectively) that transport the indicated compounds. HxnS hydroxylates nicotinic acid (NA) to 6-hydroxynicotinic acid (6-NA). HxnX operates in peroxisomes and converts 6-NA to 2,5-dihydroxypyridine (2,5-DP), which is subsequently hydroxylated by HxnV to 2,3,6-trihydroxypyridine (2,3,6-THP). HxnT and a yet-unknown alkene reductase (UE1) partially saturate the pyridine ring of 2,3,6-THP to 5,6-dihydroxypiperidine-2-one (5,6-DHPip-2-O), which is then converted to 3-hydroxypiperidine-2,6-dione (3-HPip-2,6-DO) by HxnW, a NAD-dependent polyol dehydrogenase type enzyme. The ring of 3-HPip-2,6-DO is opened by the cyclic imidase HxnM between N-C2 resulting in α-hydroxyglutaramate (α-HGA) formation. The nitrogen is salvaged by HxnN amide hydrolase and results in α-hydroxyglutarate (α-HG) formation. This reaction can also be catalyzed by other amide hydrolases (UE2). NA can be formed endogenously by the hydrolytic cleavage of amide group of nicotinamide (NAA) by a non-HxnR regulated deamidase. Cellular components as cell membrane, cytoplasm and peroxisome are shown and indicated by pictograms. Reaction in the peroxisome pictogram indicates the spatial separation of the referred catabolic step in the peroxisomes. Compound in square brackets mark a supposed intermediate that was not detected by UHPLC-HRMS method but deduced by the identified upstream and downstream metabolites. Enzyme with question mark (HxnY) supposedly works on the indicated step according to UHPLC-HRMS detected decrease of the amount of 2,5-DP and its oligomer derivatives in *hxnR*^*c*^*7 hxnVΔ hxnYΔ* strain. UE: unidentified enzyme. PI: physiological metabolite inducer of the pathway related *hxn* genes; Compounds in red letters are completely new metabolites, which have never been detected before neither in eukaryotic nor in prokaryotic organisms. (Created with BioRender.com)

We systematically deleted all *hxn* genes (*hxnS* and *hxnR* deletions were published previously (10)) in both *hxnR*^*+*^ (wild type) and *hxnR*^*c*^*7* (where the HxnR transcription factor is constitutively active) backgrounds. The resulting strains were tested for the utilization of the commercially available NA derivatives as N-sources or as inducer precursors (Fig. 3*A*). Catabolism of 6-NA in these strains was tracked by UHPLC-HRMS followed by the identification of the chemical structure of two purified metabolites by NMR (Fig. 4*A*, *Sl Appendix* Tables S1 and S2).

**Fig. 3.**
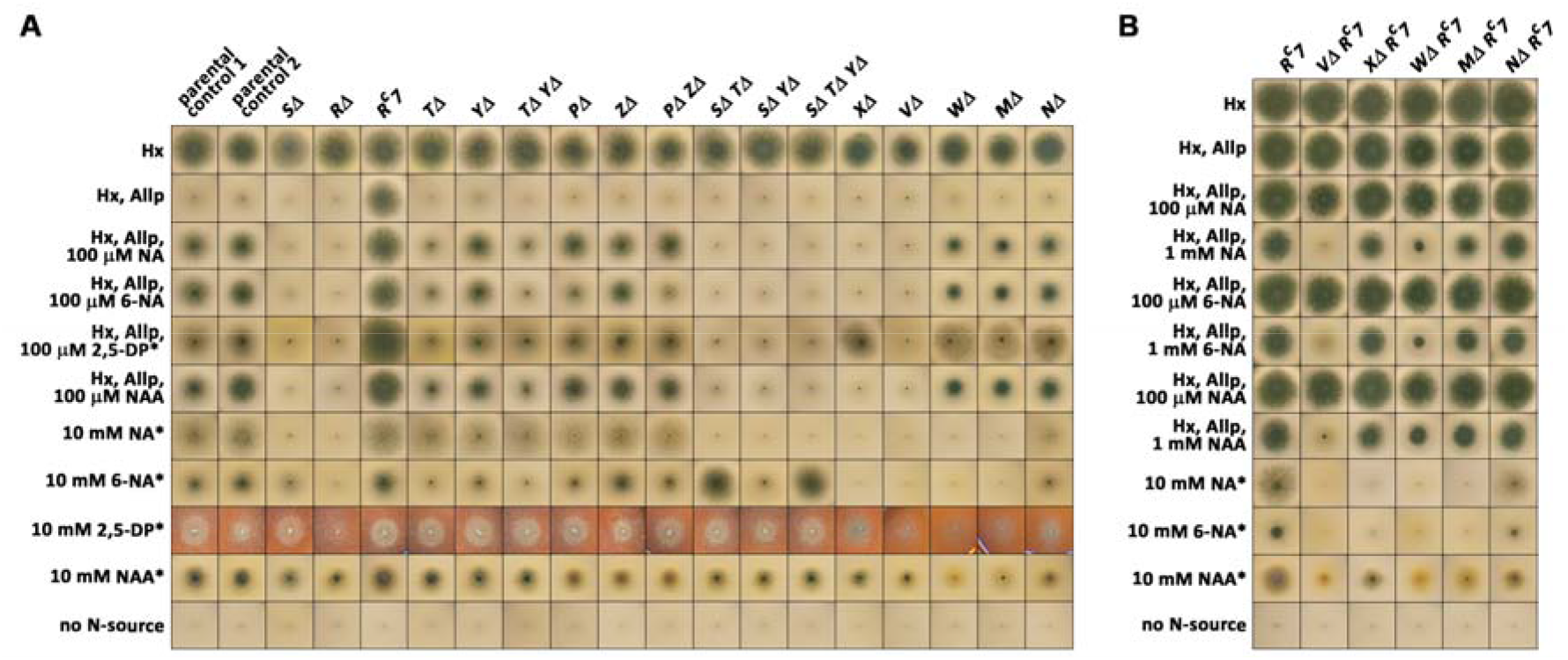
Utilization, inducer and inhibition tests of *hxn* mutants. (*A*) Utilization of different nitrogen sources by mutants described in this article in a *hxnR*^*+*^ wild type background (except for *hxnRΔ* and *hxnR*^*c*^*7* controls). (*B*) Utilization of different nitrogen sources by some *hxn* gene deletion mutants in an *hxnR*^*c*^*7* (constitutive) background. Above each column we indicate the relevant mutation carried by each tested strain. Hx indicates 1 mM hypoxanthine as the sole nitrogen source. Hx, Allp, as above including 5.5 mM allopurinol, which fully inhibits HxA but not HxnS (therefore Hx utilization depends on the activation of HxnR-regulon-belonging HxnS (for details see (10)). NA and 6-NA indicate, respectively, nicotinic acid and 6-OH nicotinic acid added as the sodium salts (see Materials and Methods section). 2,5-DP and NAA indicate, 2,5-dihydroxypyridine and nicotinamide, respectively. Other relevant concentrations are indicated in the figure. Plates were incubated for 3 days at 37 °C except those marked by asterisk (*), which were incubated for 4 days. The relevant *hxn* genes are symbolized by only the capital letter indicating the locus name. Strains used: parental control 1 (HZS.120, parent of *SΔ, TΔ, YΔ*), parental control 2 (TN02 A21, parent of *RΔ, PΔ, ZΔ, XΔ, VΔ, NΔ*) are wild type for all *hxn* genes. Mutant strains: *SΔ* (HZS.599), *RΔ* (HZS.614), *R*^*c*^*7* (FGSC A872), *TΔ* (HZS.222), *YΔ* (HZS.223), *TΔ YΔ* (HZS.502), *PΔ* (HZS.221), *ZΔ* (HZS.226), *PΔ ZΔ* (HZS.480), *SΔ TΔ* (HZS.892), *SΔ YΔ* (HZS.558), *SΔTΔ YΔ* (HZS.569), *VΔ* (HZS.294), *XΔ* (HZS.726), *WΔ* (HZS.393), *MΔ* (HZS.293), *NΔ* (HZS.288), *VΔ R*^*c*^*7* (HZS.309), *XΔ R*^*c*^*7* (HZS.310), *WΔ R*^*c*^*7* (HZS.517), *MΔ R*^*c*^*7* (HZS.308) and *NΔ R*^*c*^*7* (HZS.306). The complete genotypes are given in the *Sl Appendix* Table S3.

**Fig. 4:**
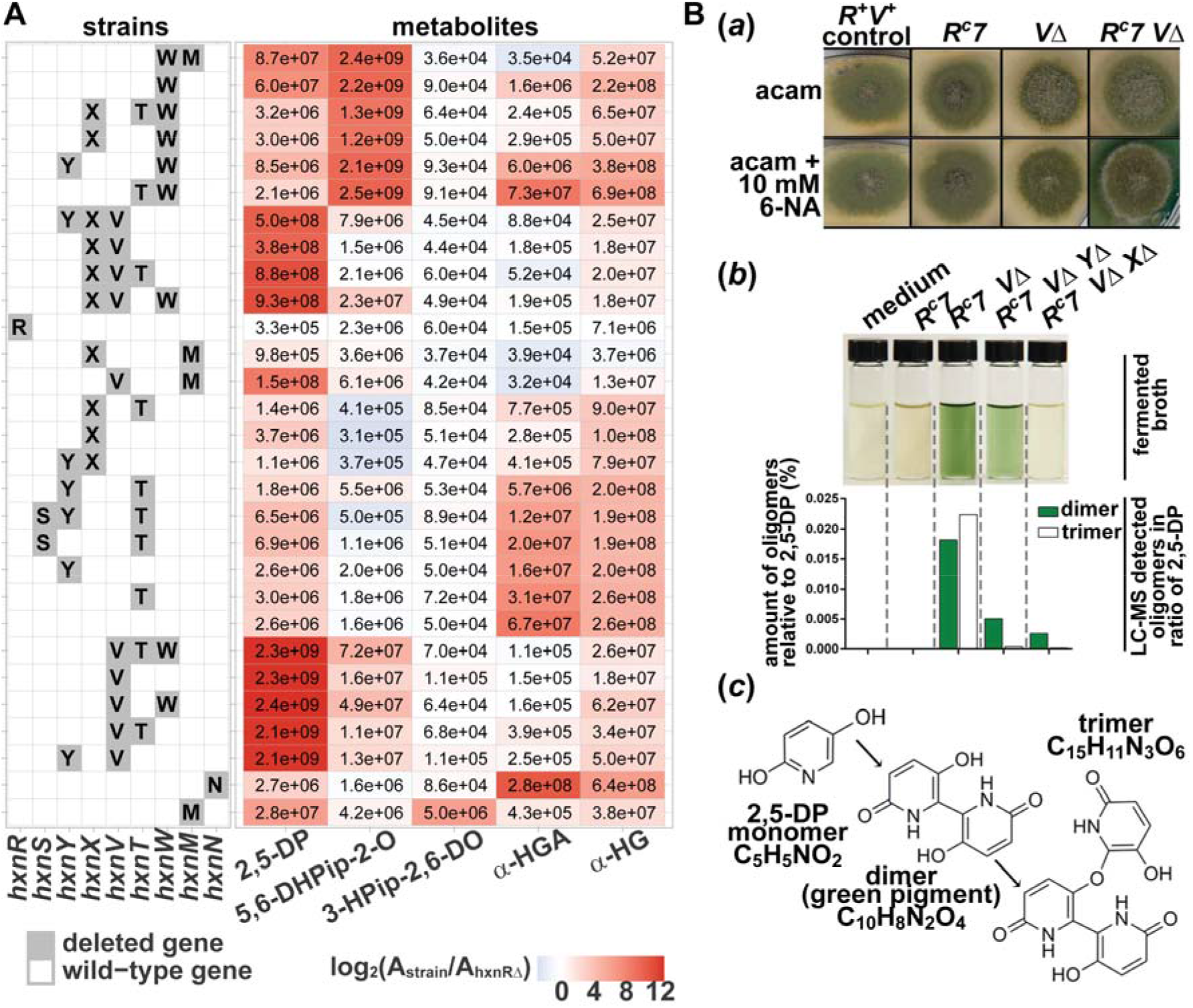
Accumulation of metabolites in various single and multiple *hxn* gene deletion mutants in a constitutive *hxnR*^*c*^*7* background. (*A*) Heat map of selected metabolites in control strains and NA catabolism impaired single and multi-deletion strains. The table to the left of this panel indicates the genotype of the used strains by a single letter code for each of the deleted *hxn* genes. The row, where no *hxn* gene deletion is indicated refers to the *hxnR*^*c*^*7* control strain. The table to the right of Panel A shows the heat map of UHPLC-HRMS measured metabolites for each strain. Numbers within the cells correspond to raw peak area values, whereas the heat map colors correspond to log_2_ fold change of peak area values relative to that of the transcription factor-deleted strain (*hxnRΔ*) (*Sl Appendix* Table S1). The data shown was obtained from mycelial extracts, except that 3-HPip-2,6-DO compound was detected and measured exclusively in the culture broth. Abbreviated compounds 2,5-DP: 2,5-dihydroxypyridine, 5,6-DHPip-2-O: 5,6-dihydroxypiperidine-2-one, 3-HPip-2,6-DO: 3-hydroxypiperidine-2,6-dione, α-HGA: α-hydroxyglutaramate, α-HG: α-hydroxyglutarate. Strains used (the relevant *hxn* genes are symbolized by only the capital letter indicating the locus name): *W*Δ *M*Δ (HZS.588), *W*Δ *R*^*c*^*7* (HZS.517), *X*Δ *T*Δ *W*Δ *R*^*c*^*7* (HZS.904), *X*Δ *W*Δ *R*^*c*^*7* (HZS.751), *Y*Δ *W*Δ *R*^*c*^*7* (HZS.898), *T*Δ *W*Δ *R*^*c*^*7* (HZS.894), *Y*Δ *X*Δ *V*Δ *R*^*c*^*7* (HZS.901), *X*Δ *V*Δ *R*^*c*^*7* (HZS.783), *X*Δ *V*Δ *T*Δ *R*^*c*^*7* (HZS.899), *X*Δ *V*Δ *W*Δ *R*^*c*^*7* (HZS.750), *R*Δ (HZS.614), *X*Δ *M*Δ (HZS.582), *V*Δ *M*Δ (HZS.584), *X*Δ *T*Δ *R*^*c*^*7* (HZS.798), *X*Δ *R*^*c*^*7* (HZS.812), *Y*Δ *X*Δ *R*^*c*^*7* (HZS.810), *Y*Δ *T*Δ *R*^*c*^*7* (HZS.903), *ST*Δ *Y*Δ *R*^*c*^*7* (HZS.912), *ST*Δ *R*^*c*^*7* (HZS.911), *Y*Δ *R*^*c*^*7* (HZS.429), *T*Δ *R*^*c*^*7* (HZS.427), *R*^*c*^*7* (FGSCA872), *V*Δ *T*Δ *W*Δ *R*^*c*^*7* (HZS.902), *V*Δ *R*^*c*^*7* (HZS.309), *V*Δ *W*Δ *R*^*c*^*7* (HZS.749), *V*Δ *T*Δ *R*^*c*^*7* (HZS.748), *Y*Δ *V*Δ *R*^*c*^*7* (HZS.747), *N*Δ *R*^*c*^*7* (HZS.306), *M*Δ *R*^*c*^*7* (HZS.308). The complete genotypes are given in the *Sl Appendix* Table S3. (*B*) Green pigment formation from 2,5-DP in a *hxnR*^*c*^*7 hxnVΔ* strain. *B(a)*: Pigment formation in solid medium. Strains were grown on MM with 10 mM acetamide (acam) as sole N-source without or with addition of 10 mM 6-NA (as the sodium salt). Strains used in this experiment: *R*^*+*^ *V*^*+*^: *hxnR*^*+*^ *hxnV*^*+*^ control (HZS.120); *R*^*c*^*7*: *hxnR*^*c*^*7* (FGSC A872); *VΔ*: *hxnVΔ* (HZS.294); *R*^*c*^*7 VΔ*: *hxnR*^*c*^*7 hxnVΔ* (HZS.309). The complete genotypes are given in the *Sl Appendix* Table S3. *B(b)*: Pigment formation in culture broth of each tested strain, compared with sterile medium. Underneath, the UHPLC-HRMS measured amounts of the dimer (green pigment) and trimer forms of 2,5-DP in the corresponding broths relative to the amount of 2,5-DP (%) are shown. Strains were grown in MM with 10 mM acetamide as sole N-source for 14 h and the mycelia were then transferred to MM without acetamide but supplemented with 10 mM 6-NA (as the sodium salt) substrate and further incubated for 24 h. Color of filtered ferment broths were photographed and subsequently analyzed by UHPLC-HRMS (see Materials and methods). Strains used in this experiment: *R*^*c*^*7* and *R*^*c*^*7 V*Δ are the same as in panel *B(a)*; *R*^*c*^*7 V*Δ *Y*Δ: *hxnR*^*c*^*7 hxnV*Δ *hxnY*Δ (HZS.747) and *R*^*c*^*7 V*Δ *X*Δ: *hxnR*^*c*^*7 hxnV*Δ *hxnX*Δ (HZS.783). *B(c)*: Deletion of *hxnV* results in the accumulation of 2,5-DP, which is non-enzymatically transformed into dimer and trimer forms. UHPLC-HRMS results for 2,5-DP are detailed in panel (*A*). Retention times of dimer and trimer forms were 1.58 and 5.63 min, and the accurate masses of precursor ions [M+H]^+^ were 221.0556 and 330.0738, respectively.

These growth tests indicate whether the tested metabolites are a nitrogen source for each strain, but also, whether in a given deletion strain the hitherto unidentified physiological inducer metabolite is synthesized or not (Fig 3*A*). To this latter end we monitor induction of *hxnS*. HxnS can catalyze the hydroxylation of hypoxanthine (Hx) to xanthine, and differently from the canonical xanthine dehydrogenase (HxA) is resistant to allopurinol (Allp) inhibition (13). Thus, if the physiological inducer metabolite is produced, a given strain would utilize Hx as a nitrogen source in the presence of Allp (Fig. 3*A*). This growth on Hx may be diminished or abolished if the accumulated pathway metabolite is toxic (Figs. 3*A* and 3*B*).

### Transporters

Two genes, *hxnP* and *hxnZ* map in cluster 1/VI and encode putative transporters of the Major Facilitator Superfamily with 12-transmembrane domains (*Sl Appendix* Figs. S1*A* and S1*C*) (11). The nearest characterized homolog of HxnP is the high-affinity nicotinate transporter TNA1 of *S. cerevisiae* (27% identity), while there is no close characterized homolog of HxnZ. Interestingly, the most likely orthologue of TNA1 in *A. nidulans* (encoded by AN5650 and sharing 31% amino acid (AA) identity with TNA1) and also its apparent paralogue in the genome (AN11116) show higher similarity with TNA1 than HxnP. While expression of AN5650 is completely independent from HxnR and NA or 6-NA induction (*Sl Appendix* S1*B*), *hxnP* shows a pattern of regulation identical to that of *hxnS* and the other enzyme-encoding genes of the clusters (10). This may signify a divergence in substrate specificity and/or a redundancy of nicotinate transporters.

Deletion of *hxnZ* impairs, but not abolishes the growth on 2,5-dihydroxypyridine (2,5-DP) and nicotinamide (NAA) as a nitrogen source and did not result in any visible impairment of growth on either NA or 6-NA (Fig. 3*A*). Deletion of *hxnP* affects very slightly the utilization of NA as nitrogen source, but clearly that of 6-NA and NAA (Fig. 3*A*). The inducer test on Hx N-source supplemented with Allp and an inducer precursor showed that a deletion of *hxnZ* slightly affects growth, while deletion of *hxnP* clearly affects the uptake of 6-NA compared to their parental control (control 2 on Fig. 3*A*). The phenotype of *hxnPΔ hxnZΔ* double mutants is identical to that of the *hxnP* single mutant (Fig. 3*A*). Deletion of either *hxnP* or *hxnZ* does not affect the nicotinate supplementation of *nicB8* auxotrophy, which can be achieved at much lower concentrations of NA (as low as 1 µM) than that those necessary for its utilization as a sole nitrogen source (10 mM) or as inducer precursor of *hxn* genes (100 µM). This is consistent with a redundancy of NA transporters in the genome and with HxnP encoding a low-affinity transporter for 6-NA, NAA and NA (Fig. 2).

### Nicotinamide utilization

One mole of N can be obtained by deamination of NAA through the action of a NAA deaminase, similar to Pnc1p in *S. cerevisiae* (14), independently from further catabolism of NA (see the growth of *hxnRΔ* and *hxnSΔ* in Fig. 3*A*). The putative NAA deaminase of *A. nidulans* encoded on Chromosome II (AN3809) is well expressed under conditions where the genes of the HxnR regulon are not expressed at all (RNA seq experiments by (15)), thus the expression of this gene must be independent of NA induction and HxnR function. The impaired utilization of NAA by *hxnWΔ, hxnMΔ* and *hxnNΔ* strains compared to *hxnRΔ*, where no *hxn* gene is expressed, is a diagnostic test of the toxicity of accumulated metabolic intermediates (Figs. 3*A* and 3*B*).

### Conversion of 6-NA to 2,5-DP occurs in the peroxisome by the 6-NA monooxygenase HxnX

Previous work has shown that HxnS catalyzes the hydroxylation of NA to 6-NA ((10) and references therein). Deletion of *hxnX* prevents the utilization of 6-NA but not 2,5-DP as a nitrogen source (Fig. 3*A*). Strains deleted for this gene are also defective in the induction of *hxnS* by 6-NA but not by 2,5-DP and an *hxnX* deletion blocks the 2,5-DP accumulation in *hxnR*^*c*^*7 hxnVΔ* mutant (Figs. 3*A*, 4*A* and 4*B*).

HxnX is a monooxygenase (Fig. 5*A*). Its closest known structural homolog is the 6-NA 3-monooxygenase, NicC (PDB code: 5eow), from *Pseudomonas putida* KT2440 (16) (Fig. 5*A*). His232 and Tyr236 residues of HxnX and their spatial orientation correspond to the 6-NA substrate binding His211 and Tyr215 residues of NicC from *P. putida* KT2440 (16) (Fig. 5*A*). The six additional AA residues, His47, Cys202, Met213, Val227, Thr228 and Gly229, which are involved in the formation of the active site (16) are not conserved in HxnX (Gln59, Val223, Val234, Val247, Leu248 and Leu249, respectively) (Fig. 5*A*). Similarly to NicC (2, 17), HxnX is proposed to require NADH, FAD and O_2_ to replace the carboxyl group with a hydroxyl group on the 6-NA substrate that results in 2,5-DP formation. HxnX includes a canonical PTS-1 peroxisome targeting signal (SRL) at its C-terminal end (*Sl Appendix* Fig. S2). An N-terminal Gfp-HxnX fusion fully complements the growth phenotype of *hxnXΔ* and co-localizes with a peroxisomal marker (Fig. 5*E*).

**Fig. 5.**
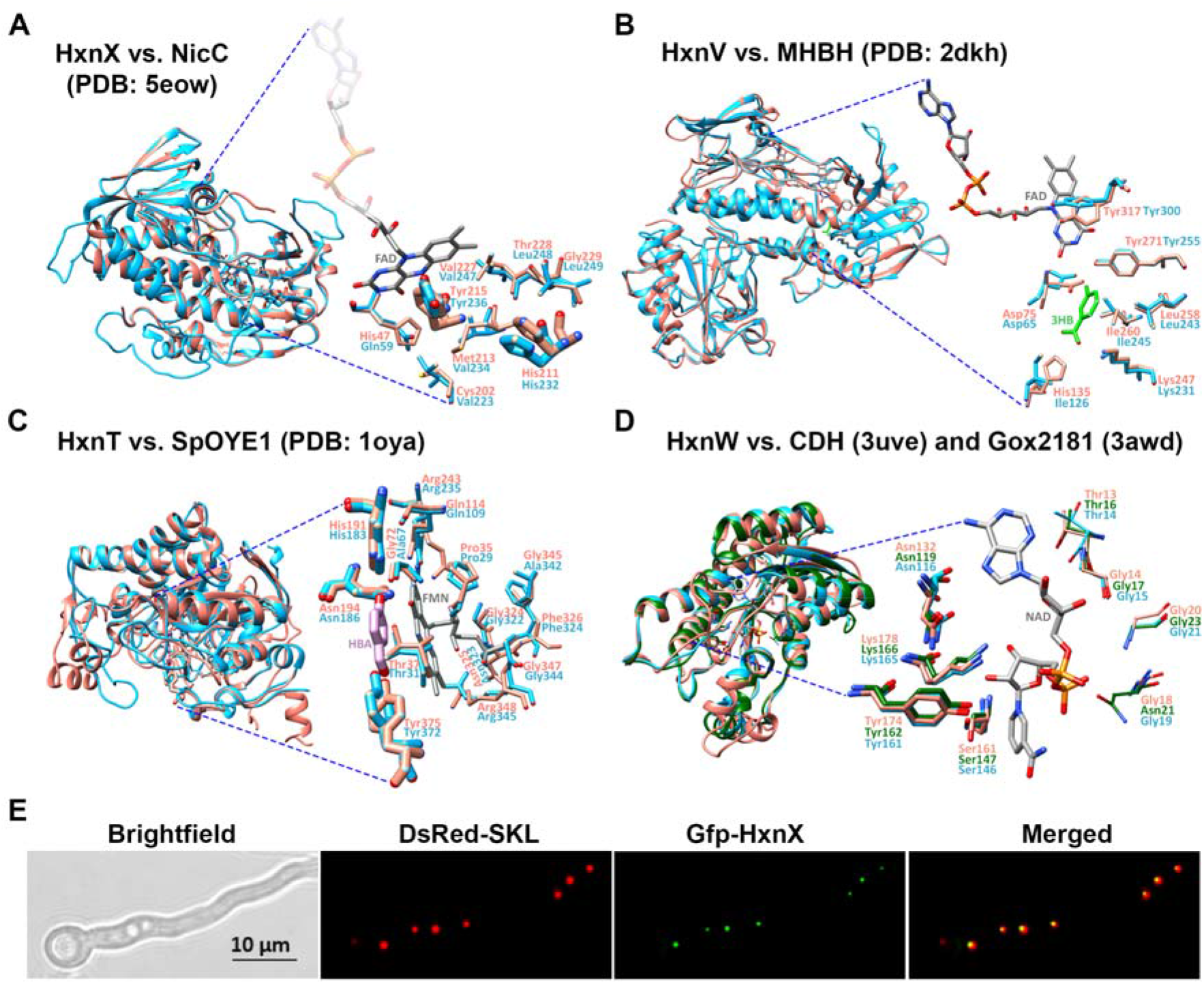
Superposition of the structural models of HxnX, HxnV, HxnT and HxnW with their closest known structural homologs. Hxn enzymes are shown in blue; the structural homologs are shown in salmon and green colors. For each model, the image to the left shows the superposition of the compared proteins in ribbon view and includes the modeled substrate and/or cofactor. The substrate interacting side chains are shown as magnified insets in stick view to the right of each model. FMN, FAD and NAD cofactors are shown by grey sticks. (*A*) HxnX *versus* 6-NA 3-monooxygenase (NicC) from *Pseudomonas putida* (PDB code: 5eow) (16). Thick sticks show 6-NA binding residues, while thin sticks show additional active site residues of NicC. (*B*) HxnV *versus* 3-hydroxybenzoate hydroxylase (MHBH) from *Comamonas testosteroni* (PDB code: 2dkh) (18). 3HB: 3-hydroxybenzoic acid (3HB) substrate (green sticks). Sticks show the 3HB substrate binding residues of MHBH. (*C*) HxnT *versus* old yellow enzyme 1 (OYE1) of *Saccharomyces pastorianus* (PDB code: 1oya) (19). HBA: para-hydroxybenzaldehyde ligand of SpOYE1 (green sticks). Thin sticks: FMN binding residues; thick sticks: HBA binding residues of SpOYE1. (*D*) HxnW *versus* the polyol dehydrogenase enzyme Gox2181 from *Gluconobacter oxidans* (PDB code: 3awd) and carveol dehydrogenase CDH from *Mycobacterium avium* (PDB code: 3uve) (20, 21). Thick sticks: active site residues in Gox2181 and CDH (20-22); thin sticks: residues of TG(X)_3_GXG NAD(P) binding motif in HxnW, characteristic of the fungal-type ketoreductases. Quality assessments of the Hxn models are summarized in *Sl Appendix* Table S4. (*E*) Subcellular localization of the Gfp-HxnX fusion protein. Gfp-HxnX is co-expressed with DsRed-SKL (peroxisome targeted red fluorescent protein (23, 24)) in strain HZS.579. Fluorescent microscopy was carried out by using Zeiss 09 and 15 filter sets for DsRed and Gfp, respectively. Conidia were germinated for 6.5 h at 37 °C on the surface coverslips submerged in MM prior to microscopy. Scale bar represents 10 μm.

The PTS-1 signal is conserved among the HxnX proteins present in other Pezizomycotina (11). No other *hxn* encoded enzyme carries a sub-cellular localization signal, which however does not exclude the possibility that the corresponding pathway-step(s) may occur in an organelle.

While constructing double mutant strains, we were surprised that, the *hxnSΔ hxnTΔ* double deletion strain utilizes 10 mM 6-NA more efficiently than the wild type control or the single *hxnSΔ* or *hxnTΔ* deletion mutants (Fig. 3*A*). The ORFs of the two divergently transcribed genes were deleted in the double mutant, the intergenic region between the start codons was left intact, excluding any cis-acting regulatory effects on other genes of the cluster (see Materials and methods). The explanation of this phenotype may relate to the intracellular pool of NAD/NADH. NAD is the final electron acceptor of HxnS (8), and the presumed electron donor of HxnT (*Sl Appendix* Fig. S3). Deletion of both cognate genes may increase the intracellular NAD/NADH pool, thus facilitating the activity of the peroxisomal HxnX, which as a monooxygenase necessitates NADH to reduce the second oxygen atom in O_2_. It seems paradoxical that the co-induction of *hxnS* and *hxnT* with *hxnX* may actually impair the utilization of 6-NA.

### Subsequent metabolism of 2,5-DP depends on the 2,5-DP monooxygenase, HxnV

N-source utilization tests showed that HxnV acts downstream of NA, 6-NA and 2,5-DP (Fig. 3*A*). Induction tests (Hx Allp rows) are completely consistent with the above, in an *hxnV*Δ strain 2,5-DP does not act as an inducer. These results place the physiological inducer of the pathway downstream form 2,5-DP. In an *hxnR*^*c*^*7* background, where all other *hxn* genes are constitutively expressed (10, 11) a *hxnVΔ* strain accumulates 2,5-DP (Fig. 4*A*) in media supplemented with 10 mM 6-NA, indicating that 2,5-DP is its substrate. This strain also secretes a green pigment (detected both visually and by UHPLC-HRMS analysis), seen both in solid medium around the colonies and in fermented broth (Fig. 4*B*). The green pigment was identified as the dimer form of 2,5-DP (Fig. 4*B*). A green pigment formation by non-enzymatic transformation of 2,5-DP was reported in the *P. putida* NicX loss-of-function mutant, blocked in the catabolism of 2,5-DP (25) and in a *P. fluorescens* strain grown on NA medium (26). The formation of the pigment is almost completely blocked in a *hxnR*^*c*^*7 hxnX*Δ *hxnV*Δ strain, consistent with the position of the HxnX protein in the pathway as the main enzyme catalyzing the formation of 2,5-DP (see above) but also diminished in an *hxnR*^*c*^*7 hxnY*Δ *hxnV*Δ strain, which suggests that HxnY may contribute to the formation of 2,5-DP from 6-NA (Figs. 2 and 4*B*).

HxnV includes a phenol 2-monooxygenase domain (PRK08294) and shows remarkable structural similarity to 3-hydroxybenzoate hydroxylase (MHBH), from *Comamonas testosteroni* (PDB code: 2dkh) (Fig. 5*B*) as well as to phenol 2-monooxygenase (PHOX) from *Trichosporon cutaneum* (PDB code: 1pn0) (Fig. 5*B* and *Sl Appendix*. Fig. S4). The phenol ring interacting residues of MHBH (Asp75, Leu258, Ile260 and Tyr271) together with their spatial orientation are fully conserved in HxnV (Asp65, Leu243, Ile245 and Tyr255), while the carboxyl group binding Lys247 and His135 residues of MHBH are partially conserved in HxnV (Lys231 and Ile126 in HxnV) (Fig. 5*B*) (18). Thus, it is not unreasonable and in agreement with data shown above that 2,5-DP be the substrate of HxnV, and by analogy between HxnV and its known structural homologs, HxnV may hydroxylate the 6-carbon of 2,5-DP resulting in 2,3,6-trihydroxypyridine (2,3,6-THP) formation (Fig. 2). This metabolite was not detected in the metabolome of any of the mutants, however the structurally identified upstream and downstream metabolites (2,5-DP and 5,6-dihydroxypiperidine-2-one (see below), respectively) suggest that 2,3,6-THP is almost certainly the product of HxnV (Figs. 2 and 4*A*).

### The 2,3,6-THP alkene reductase HxnT catalyzes the reduction of the pyridine ring

Accumulation of a saturated derivative of 2,3,6-THP, 5,6-dihydroxypiperidine-2-one (5,6-DHPip-2-O) was exclusively observed in the metabolome of an *hxnR*^*c*^*7 hxnWΔ* mutant (Fig. 4*A* and see *Sl Appendix* Table S2 for NMR results). This compound has not been detected previously in either eukaryotes or prokaryotes, and has not been synthesized chemically. 5,6-DHPip-2-O is altogether a new compound. The accumulation identified 5,6-DHPip-2-O as the substrate of HxnW but also implies that an upstream alkene reductase enzyme (HxnT, see Fig. 2 and below) acts on the hitherto undetected product of HxnV. Logically the latter has to be 2,3,6-THP. The putative alkene reductase, which supposedly converts 2,3,6-THP to the 5,6-DHPip-2-O, is HxnT (a member of the “old yellow enzymes” group). Comparison of the structural model of HxnT with its closest known structural homolog, old yellow enzyme 1 (OYE1) of *Saccharomyces pastorianus* (PDB code: 1oya) showed that the para-hydroxybenzaldehyde binding residues of SpOYE1 (His191, Asn194, Tyr375) are remarkably conserved in HxnT (His183, Asn186, Tyr372) and that the FMN binding residues are almost completely conserved in HxnT (19) (Fig. 5*C* and *Sl Appendix* Fig. S5 for further details). An *hxnT*Δ strain shows a leaky growth phenotype, most noticeably on 2,5-DP (Fig. 3*A*). The utilization of Hx in the inducer-test media is reduced but still clearly visible. Both results imply that while HxnT is responsible for the metabolism of the putative 2,5-DP metabolite to 2,3,6-THP, another, yet unidentified enzyme must be catalyzing the same step. The deletion of *hxnW* identifies 5,6-DHPip-2-O as the physiological inducer of the pathway (NA and 6-NA serve as inducer precursors in the Hx Allp test in *hxnW*Δ, but not in *hxnX*Δ and to a reduced extent in *hxnT*Δ). 2,5-DP serves as an inducer precursor in *hxnX*Δ but not in *hxnV*Δ and to reduced extent in *hxnT*Δ, which is in line with a redundantly functioning additional enzyme. While induction of a whole pathway by a metabolite such as the product of the first metabolic step has been described long ago (e.g. (27)), the pathway described in this article reports the unprecedented occurrence of concerted induction by an almost terminal metabolite.

### The 5,6-DHPip-2-one ketoreductase HxnW converts the 6-enol group of the piperidine compound to a keto group

5,6-DHPip-2-O, the product of HxnT, is the substrate of HxnW. HxnW is a short chain dehydrogenase/reductase and has a structurally conserved NADB_Rossmann fold domain (22) with a TG(X)_3_GXG (14-21 AAs) motif that is characteristic of the fungal ketoreductases (*Sl Appendix* Fig. S6). Comparison of HxnW to its closest known structural homologs, NAD(H)-dependent polyol dehydrogenase Gox2181 from *Gluconobacter oxydans* (PDB code: 3awd) and carveol dehydrogenase CDH from *Mycobacterium avium* (PDB code: 3uve) (20, 21) showed the striking conformity of the active site residues (Asn119, Ser147, Tyr162 and Lys166) (Fig. 5*D*). HxnW, similarly to its structural homologs, dehydrogenates a hydroxyl group of 5,6-DHPip-2-O resulting in the formation of 3-hydroxypiperidine-2,6-dione (3-HPip-2,6-DO) (Figs. 2 and 4*A*). Similarly to 5,6-DHPip-2-O, 3-HPip-2,6-DO is also a new natural metabolite that has not been detected previously in any organism.

The 5,6-DHPip-2-O was not accumulated in a double deleted *hxnR*^*c*^*7 hxnWΔ hxnVΔ* strain, however, it was accumulated in a *hxnR*^*c*^*7 hxnWΔ hxnXΔ* strain. This implies that a second enzyme activity may be capable of metabolizing 6-NA to 2,5-DP. We propose this enzyme to be HxnY (Fig. 2). A deletion of *hxnY* diminishes the utilization of 6-NA (Fig. 3*A*). Its contribution to the 2,5-DP pool must be minor, as an *hxnX*Δ strain does not utilize at all either NA or 6-NA as a nitrogen source (Fig 3*A*). The ancillary activity of HxnY is supported by the reduced growth on Hx Allp media in the presence of inducer precursors (Fig 3*A*). The effect of *hxnY*Δ is evident on 6-NA but not on NA. This is in line with the fact that 6-NA is a better nitrogen source than NA, and thus, the activity of HxnY may not be limiting when the organism grows on NA but may be limiting on 6-NA.

HxnY is an α-ketoglutarate-dependent dioxygenase, its closest structural homolog is the thymine-7-hydroxylase (T7H) of *Neurospora crassa* (PDB code: 5c3q) (28), which catalyzes the sequential conversion of the methyl group of thymine to a carboxyl group (28, 29). The conservation of the α-ketoglutarate and Fe^2+^ binding residues and those involved in π-π stacking and hydrophobic interactions with the pyrimidine ring of T7H are consistent with the putative activity of HxnY on 6-NA (*Sl Appendix* Fig. S7).

### The 3-HPip-2,6-DO cyclic imide hydrolase HxnM catalyzes the opening of the piperidine ring

3-HPip-2,6-DO (generated by HxnW) was accumulated exclusively, albeit in small quantity, in the fermented broth of an *hxnMΔ* strain (Fig. 4*A*). HxnM shares 74.3% identity (with 100% query coverage) with a *Candida boidinii* enzyme (OWB68015) belonging to the EC 3.5.99 enzyme class (GOterm: 0016810, hydrolase activity on non-peptide C-N bonds) and 64.2% identity (with 95.4% query coverage) with AAY98498, a cyclic imide hydrolase homolog, from *P. putida* (30) (*Sl Appendix* Fig. S8). HxnM shows striking structural similarity with its closest known structural homolog, the alleged peptidoglycan deacetylase of unknown substrate specificity HpPgdA from *Helicobacter pylori* (PDB code: 3qbu), related to cyclic imidases (31) (*Sl Appendix* Fig. S8). The closest phylogenetic relative of HpPgdA is an allantoinase (PuuE) from *Pseudomonas aeruginosa*, whose natural substrate is a small cyclic imide (31, 32). We propose that HxnM opens the ring of 3-HPip-2,6-DO between a C2 carbon and nitrogen, generating α-hydroxyglutaramate (α-HGA), a compound which was detected exclusively in the metabolome of an *hxnR*^*c*^*7 hxnNΔ* strain (Figs. 2 and 4*A*). A ring opening is a necessary step to generate NH_4_^+^ which can serve as a nitrogen source. In *A. nidulans*, uniquely among studied NA-catabolizing organisms, the generation of a piperidine ring from a pyridine ring precedes the hydrolysis of a C-N bond. In the different pathways described in bacteria, the ring opening may take place by an oxidative process in an aromatic ring (such as in *P. putida*) or in a hydrolytic process on saturated or partially saturated rings in other bacteria (*Eubacterium barkeri* and *Azorhizobium caulinodans*) (2, 3, 33) (Fig. 6).

**Fig. 6.**
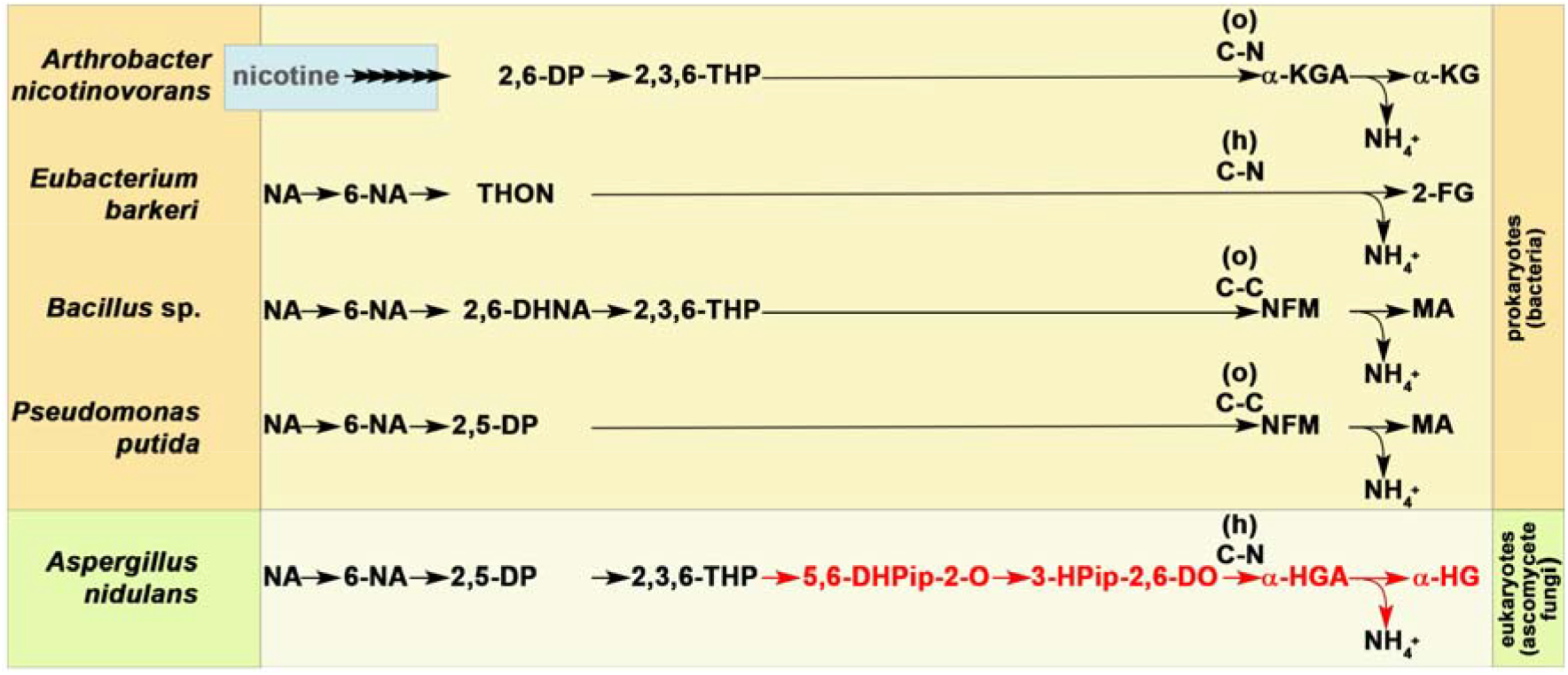
Comparative demonstration of novelties of the eukaryotic nicotinate catabolic pathway. The catabolism of nicotine by *A. nicotinovorans* involves the opening and release of the pyrrolidine ring, leading to 2,6-DP, which is further catabolized through 2,3,6-THP, an intermediate of pathways in *Bacillus* sp. as well as in *A. nidulans*. The nicotine pathway upstream to 2,6-DP is not relevant to the present work, which is indicated linked arrows and blue boxing. Red color marks those steps and pathway metabolites that have never been identified in prokaryotic NA catabolic pathways. While the eukaryotic NA catabolic pathway has only been studied experimentally in *A. nidulans*, genes encoding the whole or part of the pathway are present in many ascomycete fungi (11). Abbreviations: NA: nicotinic acid; 6-NA: 6-hydroxynicotinic acid; 2,6-DP: 2,6-dihydroxypyridine, 2,5-DP: 2,5-dihydroxypyridine; 2,6-DHNA: 2,6-dihydroxynicotinic acid; THON: 1,4,5,6-tetrahydro-6-oxonicotinic acid; 2,3,6-THP: 2,3,6-trihydroxypyridine; 5,6-DHPip-2-O: 5,6-dihydroxypiperidine-2-one; 3-HPip-2,6-DO: 3-hydroxypiperidine-2,6-dione; MA: maleamic acid; NFM: N-formylmaleamic acid; 2-FG: (*S*)-2-formylglutarate; α-KGA: α-ketoglutaramic acid; α-KG: α-ketoglutaric acid; α-HGA: α-hydroxyglutaramic acid; α-HG: α-hydroxyglutaric acid; C-C: site of ring-opening occurs between two carbons; C-N: site of ring-opening occurs between carbon and nitrogen; (o): ring-opening is oxidative; (h): ring-opening is hydrolytic.

### The α-HGA amide hydrolase HxnN is involved in nitrogen salvage from NA

HxnN is a putative amide hydrolase, its closest structural homolog is the fatty acid amide hydrolase 1 (FAAH1) from *Rattus norvergicus*. Deletion of *hxnN* diminishes but not abolishes the utilization of NA, 6-NA and 2,5-DP as sole nitrogen sources. While *hxnN* encodes the last enzyme of the *hxn* regulon, the growth tests demonstrate that (a) yet-unidentified hydrolase(s) contribute(s) to the deamidation of α-HGA (Fig. 3*A*). Several genes encoding putative paralogues of HxnN are extant in the genome of *A. nidulans* with identities to HxnN up to 39%. Superposition of the structural model of HxnN with its closest known structural homolog, FAAH1 (PDB code: 2vya), shows that the catalytic triad residues from FAAH1 involved in the hydrolysis of the amide bond, the “oxyanion hole” forming residues and the Ser residue that interacts with the catalytic triad residues (34, 35) are fully conserved in HxnN (*Sl Appendix* Fig. S8). None of the prokaryotic amide hydrolases operating in the NA catabolic routes (ω-amidases) (2, 36, 37) show considerable similarity to HxnN. Amide hydrolysis of 3-HPip-2,6-DO generates α-hydroxyglutarate (α-HG) (Figs. 2 and 4*A*), which has not been detected as intermediate in any of the elucidated prokaryotic NA catabolic routes.

### Toxicity of intermediate catabolic compounds

In an *hxnR*^*c*^*7* background all *hxn* genes are constitutively transcribed. We can thus investigate the accumulation of NA metabolites without the need for the physiological inducer metabolite of the pathway. The accumulated 2,5-DP in *hxnVΔ* is a strong inhibitor of growth, while 5,6-DHPip-2-O in *hxnWΔ* mildly, and 6-NA, 3-HPip-2,6-DO and α-HGA in *hxnXΔ, hxnMΔ* and *hxnNΔ*, respectively, slightly inhibit growth (Fig. 3*B*). Growth inhibition by pathway metabolites was also detected when acetamide was the sole N-source.

### Concluding remarks

The eukaryotic NA catabolic pathway described above shows clear differences from recently known prokaryotic pathways in steps that precede (compounds 5,6-DHPip-2-O and 3-HPip-2,6-DO) and follow (compounds α-HGA and α-HG) ring opening. Conversion of NA to 2,3,6-THP through 2,5-DP was not detected in prokaryotes, albeit these intermediates appear as elements of various pathways (Fig. 6).

E.g. 2,5-DP is formed from 6-NA in *Pseudomonas* sp., which is not hydroxylated further but the pyridine ring is cleaved between C5-C6 (2) and 2,3,6-THP in *Bacillus sp*. is formed from 2,6-dihydroxynicotinic acid (38). Notably, formation of 2,3,6-THP occurs in the nicotine catabolism by *Arthrobacter sp*. through as a product of 2,6-dihydroxypyridine metabolism (reviewed in (39)). Steps of the saturation of the pyridine ring of 2,3,6-THP to 5,6-DHPip-2-O by the OYE-related alkene reductase HxnT (and a yet-unidentified enzyme) and oxidation of 5,6-DHPip-2-O to 3-HPip-2,6-DO by the ketoreductase/polyol dehydrogenase HxnW have hitherto only been detected in this pathway (Figs. 2 and 6). Moreover, 5,6-DHPip-2-O is a completely new chemical compound. The ring opening of the piperidine ring occurs between C-N (by the cyclic imidase HxnM) generating α-HGA, which has not been found previously in NA catabolic pathways (Fig. 6). In aerobic prokaryotic pathways the ring opening occurs either between C-C of 2,5-DP (by extradiol dioxygenase) or 2,3,6-THP (in *Pseudomonas sp*. and *Bacillus sp*., respectively) or between C-N of 2,3,6-THP (in *Rhodococcus sp*. and *Arthrobacter sp*. by polyketide cyclase) generating N-formyl maleamic acid or α-ketoglutaramate (2, 36-38) (Fig. 6). In the following steps in prokaryotes, the amide is hydrolyzed by ω-amidases (2, 36, 37) not related to the HxnN amidase. The anaerobic pathway described in *E. barkeri* and *Azorhizobium caulinodans* involves the partial saturation of the pyridine ring of 6-NA that results in 1,4,5,6-tetrahydro-6-oxonicotinic acid (THON), followed by hydrolytic ring opening of THON between C-N and the simultaneous deamination (by a bifunctional enamidase in *E. barkeri*) resulting in (*S*)-2-formylglutarate formation (3, 33) (Fig. 6). While no redundantly functioning enzymes are involved in the prokaryotic routes, three steps of the fungal catabolism use alternative enzymes (HxnY and two unidentified enzymes, one functioning redundantly with HxnT, the other with HxnN) (Fig. 2). Catabolic steps downstream to 2,3,6-THP differ from those in prokaryotes and lead to the newly identified intermediate metabolites 5,6-DHPip-2-O and 3-HPip-2,6-DO (Fig. 6). The identification of these new metabolites may be of industrial or agricultural importance. The complete description of this eukaryotic pathway further illustrates the convergent evolution, both at the level of individual enzymes and at the level of a whole pathway.

## Materials and Methods

### Strains and growth conditions

The *A. nidulans* strains used in this study are listed in *Sl Appendix* Table S3. Standard genetic markers are described in http://www.fgsc.net/Aspergillus/gene_list/. Minimal media (MMs) with glucose as sole carbon source and different sole nitrogen sources were used (40, 41). The media were supplemented with vitamins (http://www.fgsc.net) according to the requirements of each auxotrophic strain. Nitrogen sources, inducers, repressors and inhibitors were used at the following concentrations: 10 mM NA or 10 mM 6-NA (1 : 100 dilution from 1 M NA or 6-NA dissolved in 1 M sodium hydroxide), 10 mM 2,5-DP added as a powder, 10 mM NAA added as a powder, 1 mM Hx added as a powder, 10 mM acetamide as sole N-sources; NA sodium salt, 6-NA sodium salt, 2,5-DP, NAA in 1 mM or 100 µM final concentration as inducers; 5.5 mM Allp as inhibitor of purine hydroxylase I (HxA) enzyme activity. Strains were grown at 37 °C for the indicated times.

For metabolite extraction, the mycelia of *hxnR*^*c*^*7* strains with different *hxn* gene deletion(s) were grown for 16 h on MM with 10 mM acetamide as sole N-source at 37 °C with 150 r.p.m. agitation, which was followed by shifting the mycelia to MM with 10 mM 6-NA as substrate without additional utilizable N-source and incubated for further 24 h.

### Gene deletions

Deletion of *hxnT/R/Y/Z/P/X/W/V/M/N* genes were constructed as described previously (42). The gene targeting substitution cassette was constructed by double-joint PCR (43), where the *riboB*^*+*^, *pabaA*^*+*^ or *pyroA*^*+*^ genes were used as transformation markers. Construction of double and triple deletion mutants or changing the *hxnR*^*+*^ genetic background of mutants to *hxnR*^*c*^*7* was carried out by standard genetic crosses or transformation followed by checking via PCR and Southern blots. DNA was prepared from *A. nidulans* as described by Specht et al. (44). Hybond-N membranes (Amersham/GE Healthcare) were used for Southern blots (45). Southern hybridizations were done by DIG DNA Labeling and Detection Kit (Roche) according to the manufacturer’s instructions. Transformations of *A. nidulans* protoplasts were performed as described by Antal et al. (46). The protoplasts were prepared from mycelia grown on cellophane (47, 48) using a 4% solution of Glucanex (Novozymes, Switzerland) in 0.7 M KCl. Transformation of 5 × 10^7^ protoplasts was carried out with 100–500 ng of fusion PCR products. Primers used in the manipulations described above are listed in *Sl Appendix* Table S5. For detailed description of single and multiple gene deletions see *Sl Appendix* Data Table S6 and Figs. S10 and S11.

### Construction and microscopy of Gfp-HxnX (N-terminal fusion) expressing strains

Construction of the *gfp*-*hxnX* expressing strain is described in details in *Sl Appendix* Data. Briefly, a bipartite cassette of the *gfp*-*hxnX* fusion was constructed by Double-Joint PCR (DJ-PCR) (43), and cloned into the pAN-HZS-1 vector (42) yielding the *gfp*-*hxnX* expression vector pAN-HZS-13, which was used to transform an *hxnXΔ* strain (HZS.534), which carries a peroxisome marker (expresses DsRed-SKL) (23, 24) (*Sl Appendix* Data Fig. S12). Transformants carrying the *gfp-hxnX* transgene from 1-10 copies were isolated. Gfp-HxnX localization was studied in HZS.579 that carried the transgene in 7 copies. Conidiospores of HZS.579 was germinated for 6.5 h on the surface of coverslips submerged in MM at 37 °C. Young hyphae were examined by fluorescence microscopy using Zeiss 09 and 15 filter sets for DsRed and GFP, respectively.

### Metabolite analysis

For metabolite extraction, 1 ml of methanol/water (8/2) was added to both 25 mg of freeze-dried mycelium and 2 ml freeze-dried fermentation broth from each cultivation followed by vortexing for 1 min and sonication at 50 W for 3 x 5 min on ice in between vortexing the samples for 30 s. After centrifugation (20 000 g, 10 min, 4 °C) the supernatants were subjected to UHPLC-HRMS analysis. UHPLC-HRMS measurements were performed using a DionexUltimate 3000 UHPLC system (Thermo Scientific) coupled to a Q Exactive Plus hybrid quadrupole-Orbitrap mass spectrometer (Thermo Scientific) operating with a heated electrospray interface (HESI). Metabolites were separated on an Acquity UPLC BEH Amide (2.1 x 100 mm, 1.7 μm) column (Waters, Hungary) thermostated at 40 °C. Acetonitrile (A) and water (B) both supplemented with 0.1% formic acid served as mobile phases. A gradient elution program was applied as follows: 0-0.5 min: 97% A, 0.5-4 min: 97-88% A, 4-10 min: 88-40% A, 10-13 min: 40% A, 13-13.5 min: 40-97% A, 13.5-27.5 min: 97% A. The flow rate was kept at 0.3 ml/min, and the injection volume was 3 µl.

All samples were analyzed in both positive and negative ionization mode using the following ion source settings: the temperature of the probe heater and ion transfer capillary, spray voltage, sheath gas flow rate, auxiliary gas flow rate and S-lens RF level were set to 300 °C, 350 °C, 3.5 kV, 40 arbitrary unit, 10 arbitrary unit and 50 arbitrary unit, respectively. For data acquisition full-scan/data-dependent MS/MS method (Full MS/ddMS2) was applied, where the full scan MS spectra were acquired at a resolution of 70,000 from m/z 50 to 500 with a maximum injection time of 100 ms. For every full scan, 5 ddMS2-scans were carried out with a resolution of 17,500 and a minimum automatic gain control target of 1.00×10^5^. Isolation window was 0.4 m/z. Instrument control and data collection were carried out using Trace Finder 4.0 (Thermo Scientific) software. The raw data files were processed by Compound Discoverer 2.1 software for chromatographic alignment, compound detection, and accurate mass determination.

All NMR experiments were accomplished on a Bruker Ultrashield 500 Plus spectrometer, solvent residual signals (methanol, DMSO) adopted as internal standards. Due to solubility issue, α-HGA was measured in it’s ammonium salt form.

### Purification of 5,6-DHPip-2-O and α-HGA

4 g and 14 g of freeze dried mycelia of 5,6-DHPip-2-O and α-HGA accumulating strains were extracted in 160 ml and 560 ml of methanol, respectively. The extracts were than evaporated to dryness and were purified with dry sample loading injection on a CombiFlash EZPrep flash chromatograph (Teledyne Isco, USA) using 0.063-0.2 mm spherical silica (Molar Chemicals, Hungary) as solid phase. The metabolite detected at *m/z* 132.0656 was separated with ethyl acetate/methanol, 4/1 (V/V) supplemented with 5% aqueous ammonia as a mobile phase resulting 5 mg material. For the metabolite detected at *m/z* 146.0461, the separation using ethyl acetate/methanol, 7/3 (V/V) supplemented with 5% aqueous ammonia was followed by additional separation step, where a mixture of methanol/water (95/5, V/V) as mobile phase was applied to achieve 6 mg purified material. At each step of the purification, the purities of the metabolites were determined via UHPLC-HRMS method described above.

### *In silico* structural analysis of Hxn proteins

Structural models of the Hxn enzymes were obtained with I-Tasser (49) followed by refining the model using ModRefiner (50) and Ramachandran plot quality assessment (results of model- and superpositioning quality assessments are summarized in *Sl Appendix* Table S4). Result of I-Tasser analysis (49) provided a list of structural homologs, those with the best C-score were chosen to superpose with the refined models.

## Acknowledgement

We thank Gábor Endre the purification of commercially available 6-hydroxynicotinic acid used as substrate in medium for metabolome analysis. A.G. was supported by grants 20391 3/2018/FEKUSTRAT, NKFIH K 123952, LP2018-15/2018. The project has received funding from the EU’s Horizon 2020 research and innovation program under grant agreement No. 739593. Work was supported by the Hungarian National Research, Development and Innovation Office (NKFIH K16-119516) and by the Hungarian Government (GINOP-2.3.2-15-2016-00012).

