## Supplementary material for "Nicotinate degradation in a microbial eukaryote: a novel, complete pathway extant in *Aspergillus nidulans*": Sl Appendix Data

### Supplementary Data

#### Deletion of the *hxnR/T/Y/Z/P/V/W/X/M/N* genes by the transformation of gene-substitution cassettes constructed with Double-Joint PCR.

Deletion of the *hxn* cluster genes was carried out as described previously (1). The tripartite transformation cassettes (composed from ("A", "B" and "C" components) were constructed with Double-Joint PCR (DJ-PCR) (2). While the "A" and "C" components drive homologous recombination, the "B" component served as selection marker. The *riboB* and/or *pabaA* and/or *pyroA* from *A. nidulans* or the *riboB* orthologue from *A. fumigatus* (Afu1g13300) were used as selection marker genes for gene-replacements. For *hxnY/P/Z/V/W/X/M* and *hxnN* deletion the *riboB* gene from *A. nidulans* was used*;* for the *hxnR* deletion the *riboB* gene from *A. fumigatus* was used; for the *hxnX*, *hxnT* deletions and the *hxnS-hxnT* double deletion, the *pabaA* gene from *A. nidulans* was used; for the *hxnZ* deletion, the *pyroA* gene from *A. nidulans* was used as the selection marker gene. Construction of the gene-replacement cassettes involved the amplification of the upstream and downstream flanking regions ("A" and "C" components of the cassette, respectively) of the targeted genes by using specific primers and the amplification of the selection marker gene ("B" component) by using chimeric primers. The 5' moieties of the forward chimera primers were specific for the 3' end of the "A" components; the 5' moieties of the reverse chimera primers were specific for the 5' end of the "C" components. Assembly of the "A", "B" and "C" components were carried out in a PCR reaction where "A"-component-specific nested forward and "C"-component-specific nested reverse primers were used. All the primers used and the sizes of PCR products for the construction of each gene-replacement cassettes are described below, in Table S6. The recipient strains used for transformation, the result of PCR-based pre-selection (by using targeted gene-specific primer pairs) of transformants, and the Southern hybridization strategy for the analysis of the transformants are also listed in Table S6. The Southern blot hybridization was carried out on restriction-enzyme-digested total DNA extracts of transformant strains and wild-type or recipient control strains with "A" or "C" component derived DNA probes using DIG-DNA labeling and detection kit (Roche) (details are shown in Table S6 and Fig. S10). Those transformant strains, which underwent the expected gene-replacement and were free from ectopic integration of the gene-replacement cassette, were used for nicotinate utilization tests (selected transformants are marked on Table S6 and Fig. S10). The co-segregation of the selection marker genes with the single deletions were tested for each selected deletion strains by analysis of the progeny of genetic crosses. The genetic crosses carried out are listed in Table S6.

**Table S6.** Detailed summary of construction and checking of gene deletion strains.

| **Targeted gene** | **Construction of gene substitution casettes by DJ-PCR** | | | | | | **Southern analysis strategy** | | | | | **Selected transformant strains** | **Co-segregation tests by genetic crosses** |
|---|---|---|---|---|---|---|---|---|---|---|---|---|---|
| | forward + reverse primers[1] and product size | | | | Recipient strain[1] | Primers for PCR tests[2] | Restriction enzymes[3] | Primers[1] for DNA probes | Controll size | Gene replacement size | Corres-ponding image* | HZS collection number | Strains used for crossing |
| | "A" component | "B" component | "C" component | Assembly of "A", "B", "C" | | | | | | | | | |
| *hxnP*[4] | 1+2 (2,435 bp) | 3+4 (2,264 bp) | 5+6 (2,438 bp) | 7+8 (5,878 bp) | TN02 A21 | 108+109 | EcoRI | 1+2 | 5,752 bp | 3,240 bp | *A* | 221 | HZS.399, CS2638 |
| *hxnY*[4] | 9+10 (2,517 bp) | 11+12 (2,266 bp) | 13+14 (2,566 bp) | 15+16 (6,396 bp) | HZS.120 | 112+113 | BamHI | 9+10 | 13,837 bp | 4,415 bp | *B* | 223 | HZS.395 |
| *hxnZ*[4] | 17+18 (2,532 bp) | 19+20 (2,266 bp) | 21+22 (2,530 bp) | 23+24 (6,584 bp) | TN02 A21 | 114+115 | BamHI | 17+18 | 13,887 bp | 5,904 bp | *C* | 226 | HZS.399, CS2638 |
| *hxnT*[5] | 28+29 (2,666 bp) | 30+31 (3,905 bp) | 32+33 (2,645 bp) | 34+35 (7,535 bp) | HZS.120 | 110+111 | EcoRI | 32+33 | 16,703 bp | 3,568 bp | *D* | 222 | HZS.397 |
| *hxnT*[6] | 40+41 (2,219 bp) | 42+43 (2,208 bp) | 44+33 (2,748 bp) | 45+35 (6,260 bp) | HZS.599 | 110+111 | EcoRI | 44+33 | 5,340 bp | 3,560 bp | *E* | 892 | - |
| *hxnR*[7] | 46+47 (3,204 bp) | 48+49 (2,607 bp) | 50+51 (2,787 bp) | 52+53 (7,611 bp) | TN02 A21 | 106+107 | XbaI | 54+55 | 8,557 bp | 5,729 bp | *F* | 614 | HZS.120 |
| *hxnX*[4] | 56+57 (2,849 bp) | 58+59 (2,208 bp) | 62+63 (2,551 bp) | 64+65 (6,861 bp) | HZS.563 | 116+117 | SacI | 56+57 | 6,195 bp | 10,055 bp | *G* | 726 | HZS.222, HZS.404, HZS.429, HZS.537 |
| *hxnX*[5] | 56+57 (2,849 bp) | 60+61 (3,846 bp) | 62+63 (2,551 bp) | 64+65 (8,499 bp) | HZS.563 | 116+117 | SacI | 56+57 | 7,379 bp | 10,055 bp | *G* | 727 | HZS.222, HZS.404, HZS.429, HZS.537 |
| *hxnV*[4] | 74+75 (2,030 bp) | 76+77 (2,208 bp) | 78+79 (2,406 bp) | 80+81 (5,202 bp) | TN02 A21 | 120+121 | EcoRI | 74+75 | 4,342 bp | 2,432 bp | *H* | 294 | FGSCA872, HZS.292 |
| *hxnW*[4] | 66+67 (2,662 bp) | 68+69 (2,208 bp) | 70+71 (3,953 bp) | 72+73 (5,948 bp) | HZS.267 | 118+119 | XbaI | 66+67 | 8,821 bp | 4014 bp | *I* | 393 | HZS.123, HZS.292 |
| *hxnM*[4] | 82+83 (3,284 bp) | 84+85 (2,208 bp) | 86+87 (3,328 bp) | 88+89 (6,018 bp) | HZS.251, HZS.267 | 122+123 | EcoRI-HindIII | 82+83 | 2,744 bp | 3,040 bp | *J* | 292, 293 | FGSCA872, HZS.297 |

| | | | | | | | | | | | | | |
|---|---|---|---|---|---|---|---|---|---|---|---|---|---|
| *hxnN*[4] | 90+91 (3,202 bp) | 92+93 (2,208 bp) | 94+95 (2,666 bp) | 96+97 (5,841 bp) | TN02 A21 | 124+125 | EcoRI-XbaI | 90+91 | 4,045 bp | 3,437 bp | *K* | 288 | FGSCA872 |
| *hxnST*[8] | 36+37 (2,825 bp) | 38+31 (3,898 bp) | 32+33 (2,645 bp) | 39+35 (7,939 bp) | HZS.564 | 126+127 | KpnI-HindIII | 39+37 | 5,336 bp | 6,500 bp | *L* | 568 | HZS.223, HZS.537, HZS.623 |
| *hxnZ*[9] | 17+18 (2,532 bp) | 25+26 (2,410 bp) | 21+22 (2,530 bp) | 27+24 (6,478 bp) | HZS.221 | 114+115 | EcoRV | 21+22 | 6,207 bp | 6,910 bp | *M* | 480 | - |
| *hxnWV*[10] | 74+75 (2,030 bp) | 76+69 (2,249 bp) | 70+71 (3,953 bp) | 80+73 (5787 bp) | HZS.404 | 128+129 130+131 | XhoI | 118+119 120+121 | 5,288 bp | - | *N* | 749 | - |
| *hxnXW*[11] | 66+67 (2,662 bp) | 68+59 (2208 bp) | 62+63 (2,551 bp) | 128+71 (6,015 bp) | HZS.404 | 130+131 116+117 | EcoRV | 116+117 118+119 | 3,546 bp | - | *O* | 751 | - |
| *hxnXWV*[12] | 74+75 (2,030 bp) | 76+59 (2,255 bp) | 62+63 (2,551 bp) | 80+73 (10,347 bp) | HZS.404 | 128+129 130+131 116+117 | XbaI | 116+117 118+119 120+121 | 10,347 bp | - | *P* | 750 | - |

[*] Indicates the corresponding Southern blot images (from panel *A*-*P*) on Fig. S10

[1] Recipient strain used for transformation

[2] Primers used for PCR are listed in *SI Appendix* Table S5.

[3] Restriction enzymes used for the digestion of total DNA

[4] The selection marker used for gene-replacement was *riboB*$^+$

[5] The selection marker used for gene-replacement was *pabaA*$^+$

[6] The *hxnT* gene was deleted in a *hxnSΔ::pabaA*$^+$ recipient strain. The selection marker gene was *riboB*$^+$ from *A. nidulans*. The shared promoter between *hxnS* and *hxnT* remained intact in the developed *hxnSΔ::pabaA*$^+$ *hxnTΔ::riboB*$^+$ double deletion strain.

[7] The selection marker used for gene-replacement was *riboB*$^+$ from *A. fumigatus.*

[8] The gene-replacement cassette targeted both of the neighboring genes, *hxnS* and *hxnT* and the shared promoter region between them. The selection marker gene was *pabaA*$^+$ from *A. nidulans*. The gene replacement deleted *hxnS* and *hxnT* simultaneously.

[9] The *hxnZ* gene was deleted in a *hxnPΔ::riboB*$^+$ recipient strain. The selection marker gene was *pyroA*$^+$ from *A. nidulans*.

[10] The gene-replacement cassette targeted both of the neighboring genes, *hxnV* and *hxnW*. The selection marker gene was *riboB*$^+$ from *A. nidulans*. The gene replacement deleted *hxnV* and *hxnW* simultaneously.

[11] The gene-replacement cassette targeted both of the neighboring genes, *hxnX* and *hxnW*. The selection marker gene was *riboB*$^+$ from *A. nidulans*. The gene replacement deleted *hxnX* and *hxnW* simultaneously.

[12] The gene-replacement cassette targeted the three neighboring genes, *hxnX, hxnW* and *hxnV*. The selection marker gene was *riboB*$^+$ from *A. nidulans*. The gene replacement deleted *hxnX*, *hxnW* and *hxnV* simultaneously.

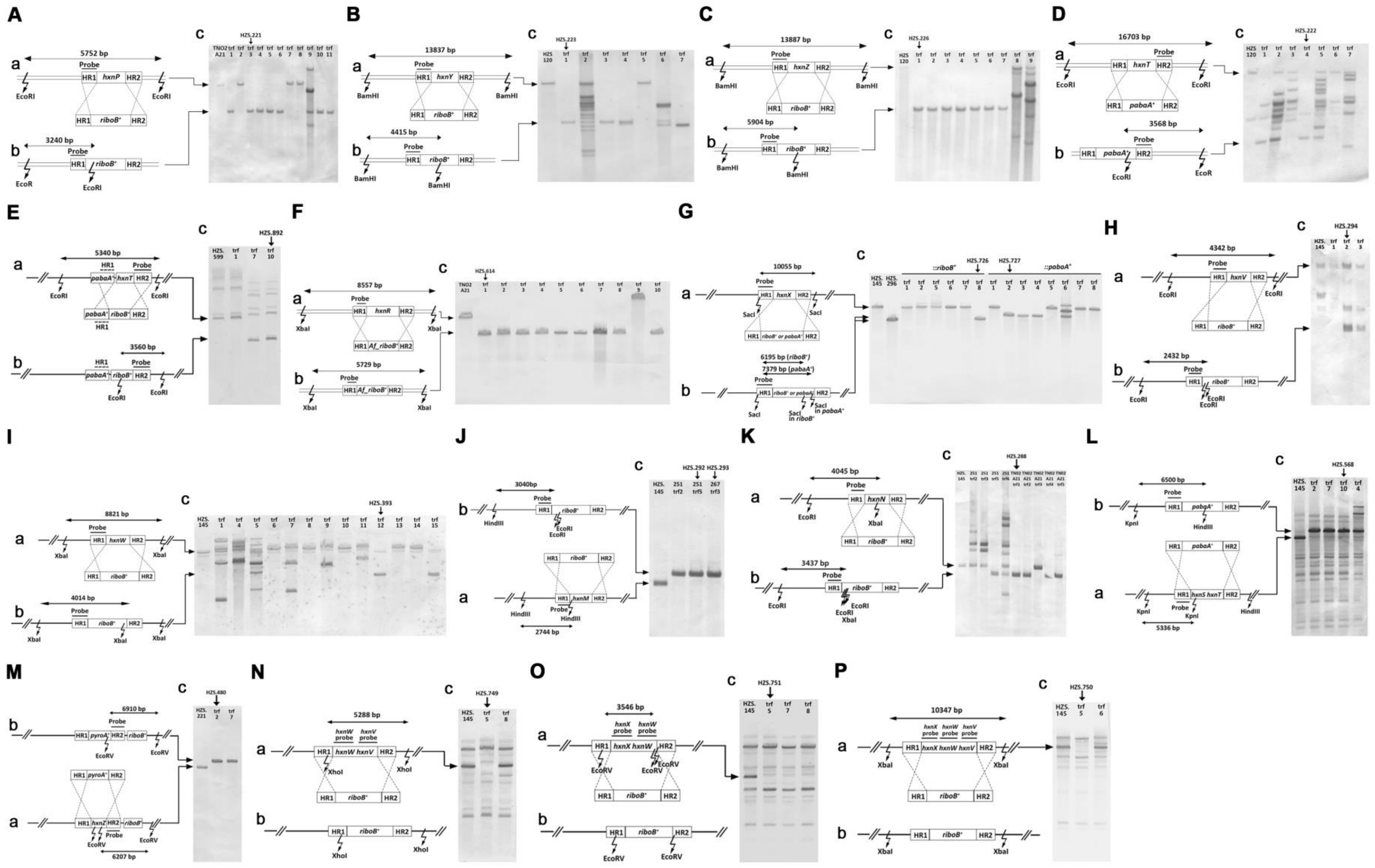

A
5752 bp
Probe
HR1 hxnP HR2
EcoRI
HR1 riboB+ HR2
3240 bp
HR1 riboB+ HR2
EcoR
EcoRI
HZS.221
TNO2 A21
B
13837 bp
HR1 hxnY HR2
BamHI
HR1 riboB+ HR2
4415 bp
HZS.223
HZS 120
C
13887 bp
HR1 hxnZ HR2
BamHI
5904 bp
HZS.226
D
16703 bp
HR1 hxnT HR2
HR1 pabaA+ HR2
3568 bp
HZS.222
E
5340 bp
HR1 pabaA+ hxnT HR2
pabaA+ riboB+ HR2
3560 bp
HZS.892
HZS. 599
F
8557 bp
HR1 hxnR HR2
XbaI
HR1 Af_riboB+ HR2
5729 bp
HZS.614
G
10055 bp
HR1 hxnX HR2
SacI
6195 bp (riboB+)
7379 bp (pabaA+)
SacI in pabaA+
SacI in riboB+
::riboB+
::pabaA+
HZS.726
HZS.727
HZS. 145
HZS. 296
H
4342 bp
HR1 hxnV HR2
2432 bp
HZS.294
I
8821 bp
HR1 hxnW HR2
4014 bp
HZS.393
J
3040bp
HindIII
HR1 hxnM HR2
2744 bp
HZS.292
HZS.293
K
4045 bp
HR1 hxnN HR2
3437 bp
HZS.288
L
6500 bp
HR1 pabaA+ HR2
KpnI
HR1 hxnS hxnT HR2
5336 bp
HZS.568
M
6910 bp
HR1 pyroA+ HR2 riboB+
EcoRV
HR1 hxnZ HR2 riboB+
6207 bp
HZS.480
HZS. 221
N
5288 bp
hxnW probe
hxnV probe
HR1 hxnW hxnV HR2
XhoI
HZS.749
O
3546 bp
hxnX probe
hxnW probe
HR1 hxnX hxnW HR2
EcoRV
HZS.751
P
10347 bp
hxnX probe
hxnW probe
hxnV probe
HR1 hxnX hxnW hxnV HR2
XbaI
HZS.750

**Fig. S10. Southern blot analysis of gene-deletions in transformed strains.**

Panels *A-P* show the Southern blot results of transformant strains (images marked with *c*), the transformation cassettes constructed by using the DJ-PCR method together with the genomic layout of the targeted genes and their flanking regions in the recipient strain (drawings marked with *a*) and the layouts of the genomic areas after the gene-replacement events (drawings marked with *b*). Targeted gene(s) and the selection marker genes used for gene-replacements are marked in drawings by *a*. The crossing overs between the HR1 and HR2 regions (homologous recombination region 1 and 2) are indicated by crossing dashed lines. HR1 is the region that was used for homologous recombination upstream to the deletion site (corresponds to component "A" in the tripartite gene-replacement cassette); HR2 is the region that was used for homologous recombination downstream to the deletion site (corresponds to component "C" in the tripartite gene-replacement cassette). Zig-zag arrows show the positions of the cleavage sites of the restriction endonucleases used for the digestion of the total DNAs. The genomic regions used as DNA probes are shown above the corresponding genomic regions. The sizes of the hybridizing control and gene-replaced fragments are schematized above the corresponding regions by double-headed arrows. The *c* images show the Southern hybridization results (filter used:Hybond N from Amersham) where the first lanes show the control strains and the following lanes show the transformant strains (trf). Vertical arrows indicate the transformants, which were selected for further experiments. The cognate strain names are indicated above each vertical arrow, while their complete genotype is listed in *Sl Appendix* Table S3. Panels *A-P* refer to Southern blot data of gene replacements listed in Table S6.

**Introduction of the dominant *hxnR$^c$7* allele *in trans* in the *hxnR$^+$ hxnSΔ-hxnTΔ* double deletion and *hxnR$^+$ hxnSΔ-hxnTΔ-hxnYΔ* triple deletion strains.**

The plasmid pAN52-1 (3) was used to construct pAN-HZS-14 vector, from which the transformation vector, pAN-HZS-17 was constructed (Fig. S11*A* and *B*, respectively). The pAN-HZS-14 vector was constructed by cloning the PCR product of the coding sequence of the *pyroA$^+$* gene from wild-type *A. nidulans* (HZS.145) with its native promoter and termination sequence (by using the 132 and 133 primers) into the HindIII site of pAN52-1 (3) (Fig. S11*A*). The *hxnR$^c$7* allele with its native promoter and termination sequence was amplified from FGSCA872 (by using the 134 and 135 primers) and cloned into the NheI-NotI sites of the pAN-HZS-14 vector. The NheI-NotI cleavage of the vector resulted in the complete elimination of the Gfp protein coding sequence from the vector and truncation of the $P_{gpdA}$ promoter. The sequence of the cloned *hxnR$^c$7* allele was checked by sequencing the cloned region (with the primers from number 148 to 151). The resulting vector was named pAN-HZS-17 (Fig. S11*B*). pAN-HZS-17 was transformed into the HZS.568 (*hxnSΔ-hxnTΔ*) and HZS.569 (*hxnSΔ-hxnTΔ-hxnYΔ*) recipient strains followed by the isolation of pyridoxine prototroph transformants. Integration of the vector was checked by PCR using primers 136 and 137 and the copy number of the integrated *hxnR$^c$7* construct was determined by qPCR using *actA* as a reference gene. The primer pairs used for qPCR were 162-163 and 154-155. The strains selected strains for further experiments were named HZS.911 (*hxnR$^c$7 hxnSΔ-hxnTΔ*) and HZS.912 (*hxnR$^c$7 hxnSΔ-hxnTΔ-hxnYΔ*) and they carried the integrated *hxnR$^c$7* vector in two and one copies, respectively.

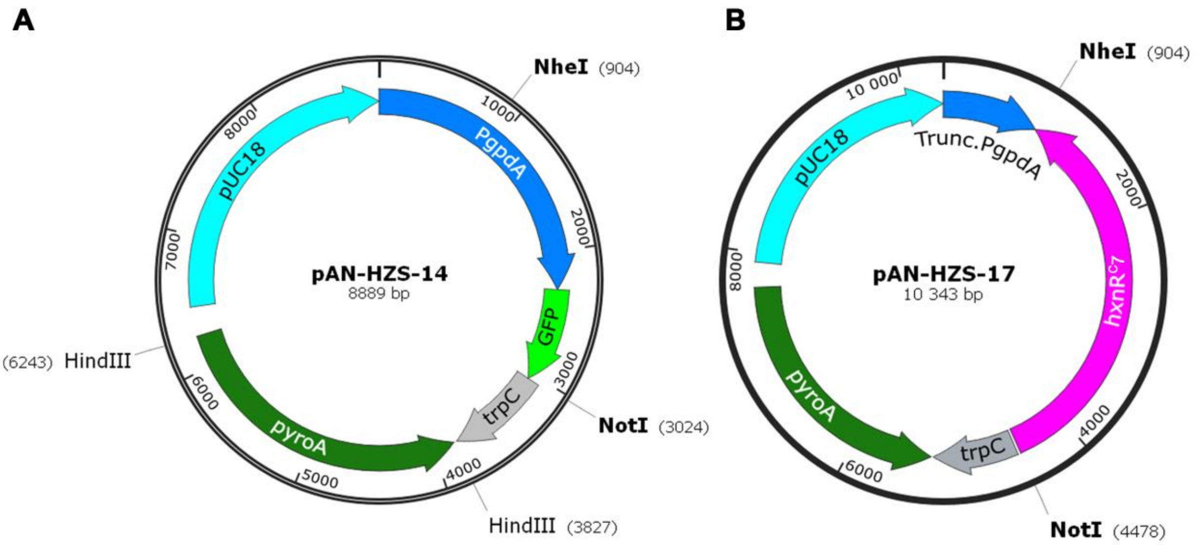


**Fig. S11. Schematic presentation of the original and resulting transformation vectors used for the construction of the *hxnR$^c$7* expression strains.** (*A*) Original vector pAN-HZS-14 used for the construction of the transformation vector. (*B*) Transformation vector pAN-HZS-17. Relevant unique restriction motifs used for cloning are shown (the numbers in parenthesis show the distance from the numbering start point in base pair units). Colored arrows show relevant components of the vectors (arrowheads indicate orientation). pUC18: standard *E. coli* vector; PgpdA: constitutive promoter of *gpdA* (glyceraldehyde-3-phosphate dehydrogenase coding gene) from *A. nidulans*; GFP: coding gene of Gfp (green fluorescence protein); trpC: termination sequence of *trpC* gene (tryptophan biosynthesis gene) from *A. nidulans*; pyroA: wild-type *pyroA* gene (coding for pyridoxine biosynthesis gene) from *A.*

*nidulans*, which serves as selection marker gene for transformation; Trunc.PgpdA: truncated *gpdA* promoter; hxnR$^{c}$7: constitutive allele of *hxnR* from *A. nidulans*. Names and sizes of the vectors are shown within the circular schemes.

**Construction of Gfp-HxnX expressing strains**

Coding sequence of Gfp (green fluorescent protein) was fused to the 5' end of *hxnX* by Double-Joint PCR (DJ-PCR) (2). *gfp* was amplified from a pAN-HZS-1 (Fig. S12*A*) template using the 138 and 139 primers. The reverse primer 139 was specific to the 3'end of *gfp* (excluding the stop codon) and carried a 24 bp long linker sequence at the 5'end that encoded 8 AAs (amino acids) (LIDTVDLD). *hxnX* was amplified from a wild-type template (HZS.145) using the 140 and 73 primers. The forward primer 140 was specific to the 5' terminus of *hxnX* (start codon included) and carried the 8 AAs linker-coding sequence at the 5' terminus. The amplified *gfp* and *hxnX* PCR products were combined into a single molecule by DJ-PCR using nested forward and nested reverse primers carrying NcoI and NotI and motifs at their 5' ends, respectively (141 and 142). The resulted *gfp-hxnX* fusion PCR product was cloned into an NcoI-NotI digested pAN-HZS-1 (Fig. S12*A*) (upon cleavage, the *gfp* gene had been eliminated from the original vector). The resulting vector (named as pAN-HZS-13) expressed the *gfp-hxnX* fusion from the constitutive *gpdA* promoter and carried the *pantoB*$^{+}$ gene that served as selection marker gene for transformation (Fig. S12*B*). pAN-HZS-13 was transformed into a peroxisome labeled (DsRed-SKL expressing) (4, 5) *hxnX*$\Delta$ strain (HZS.534). Transformants carrying the *gfp-hxnX* transgene from 1-10 copies were isolated. Copy number of *gfp-hxnX* transgene was determined by qPCR carried out with *hxnX* specific 156 and 157 and *actA* specific 162 and 163 primer pairs. The copy number of the *gfp-hxnX* transgene varied from 1-10 in the different transformants. All strains with *gfp-hxnX* transgene showed co-localization of green fluorescence with peroxisome-specific red fluorescence. Since the intensity of Gfp fluorescence was very low in strains with only 1 transgene compared to the intensity of DsRed fluorescence, we selected the strain with 7 copies of transgene (HZS.579) for fluorescence microscopy.

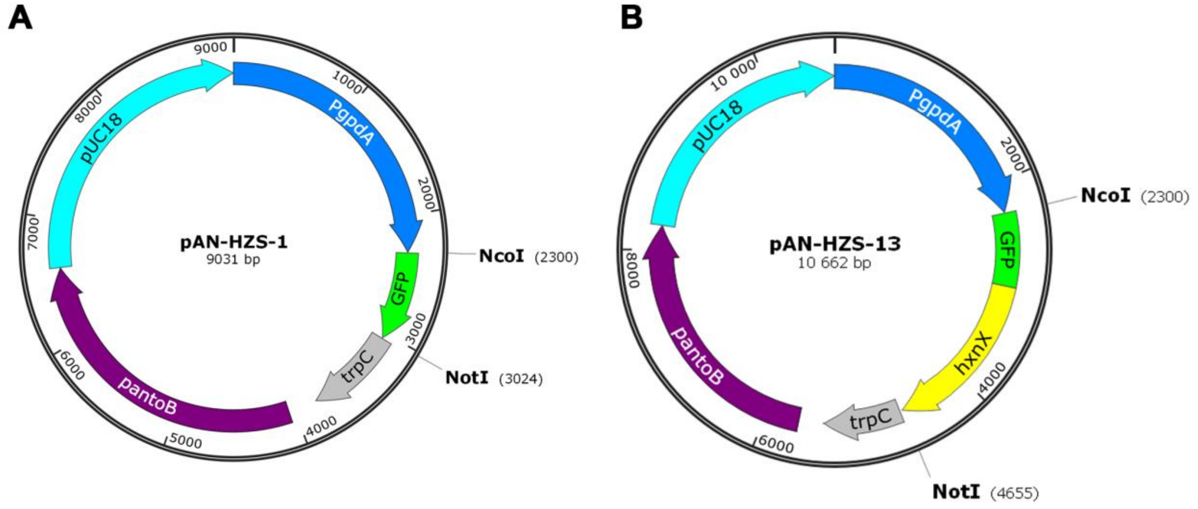


**Fig. S12. Schematic representation of the original and resulting transformation vectors used for the construction of the *gfp-hxnX* expression strains.** (*A*) Original vector pAN-HZS-1 used for the construction of the transformation vector (1). (*B*) Transformation vector pAN-HZS-13. Relevant unique restriction motifs used for cloning are shown (the numbers in parenthesis show the distance from the numbering start point in base pair units). Colored arrows show relevant components of the vectors (arrowheads indicate orientation). pUC18: standard *E. coli* vector; PgpdA: constitutive promoter of *gpdA* (glyceraldehyde-3-phosphate dehydrogenase encoding gene) from *A. nidulans*; GFP: coding gene of Gfp (green fluorescence protein); trpC: termination sequence of *trpC* gene (tryptophan biosynthesis gene) from *A. nidulans*; pantoB: wild-type *pantoB* gene (coding for a pantothenic acid biosynthesis gene) from *A. nidulans* serving as a selection marker for transformation; gfp-hxnX: *gfp* fused *hxnX* gene carrying a 8 AA coding linker sequence between *gfp* and *hxnX*. Names and sizes of the vectors are shown within the circular schemes.
