## Supplementary material for "Nicotinate degradation in a microbial eukaryote: a novel, complete pathway extant in *Aspergillus nidulans*": Sl Appendix Fig

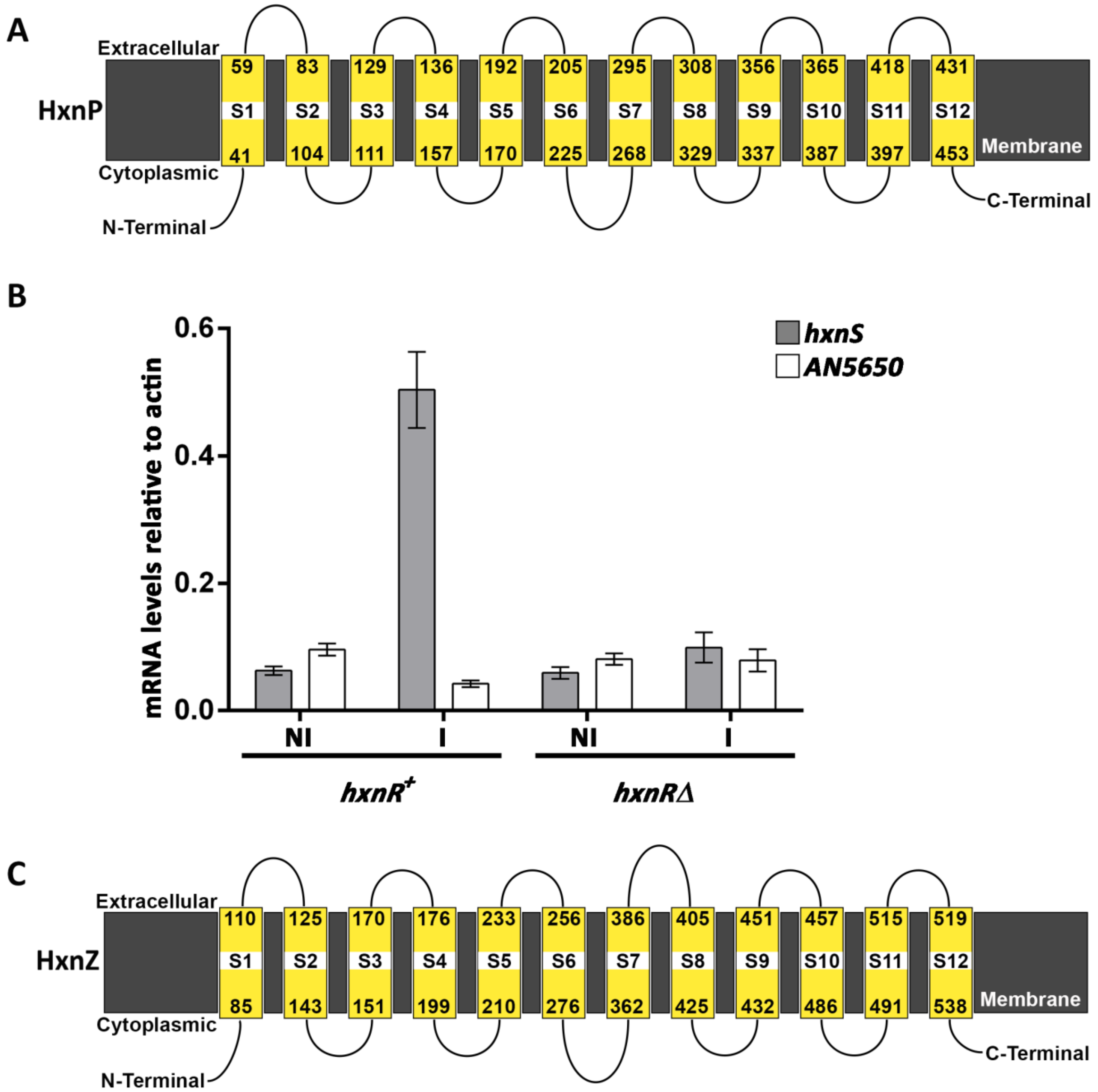


**Fig. S1. Transmembrane topologies of the HxnP and HxnZ transporters and expression profile of AN5650, the closest homolog of the yeast nicotinate transporter TNA1 in *A. nidulans*.**

(*A*) Secondary structure of HxnP obtained with Phyre2 (1). HxnP is predicted to be a 12-segment transmembrane protein of the Major Facilitator Superfamily (PF07690.13). The nearest characterized homolog of HxnP is TNA1 (shares 24% identity with HxnP), the NA transporter of *S. cerevisiae* (2). However, the most likely orthologue of TNA1 in *A. nidulans* (encoded by AN5650 and shares 31% identity with TNA1) and also its paralogues in the genome show higher similarity with TNA1 than HxnP. This may signify a divergence in substrate specificity and/or a redundancy of nicotinate transporters. While *hxnP* shows a pattern of regulation identical to that of *hxnS* and the other enzyme-encoding genes of the clusters (3), expression of AN5650 is completely independent from HxnR and NA or 6-NA induction (see panel *B*). Additionally, RNAseq data (4, 5) indicates that AN5650 is equally expressed on complete medium as in conditions of nitrogen starvation, which confirms that AN5650 is not related to NA utilization.

(*B*) Gene expression analysis of AN5650. The mRNA levels were measured by RT-qPCR and data were processed according to the relative standard curve method (6) with γ-actin (*actA*) as reference mRNA. Mycelia were grown on 1 mM acetamide as sole N-source for 8 hours at 37 °C. They were either maintained on the same media for a further 2 hours (non-induced, NI) or induced with 1 mM NA (as the sodium salt, I). Used strains were *hxnR*$^{+}$ (FGSC A26) and *hxnR*Δ (HZS.614). Standard errors of three independent experiments are shown in all RT-qPCR. Primers 152-153 (for *actA*), 158-159 (for *hxnS*) and 160-161 (for AN5650) are listed in *Sl Appendix* Table S5.

(*C*) Secondary structure of HxnZ obtained with Phyre2 (1). HxnZ is a predicted transporter of the MFS_1 superfamily with 12-segment transmembrane domain. Its closest characterized homolog in *S. cerevisiae* (17% identity) is PHO84, a high-affinity phosphate transporter.

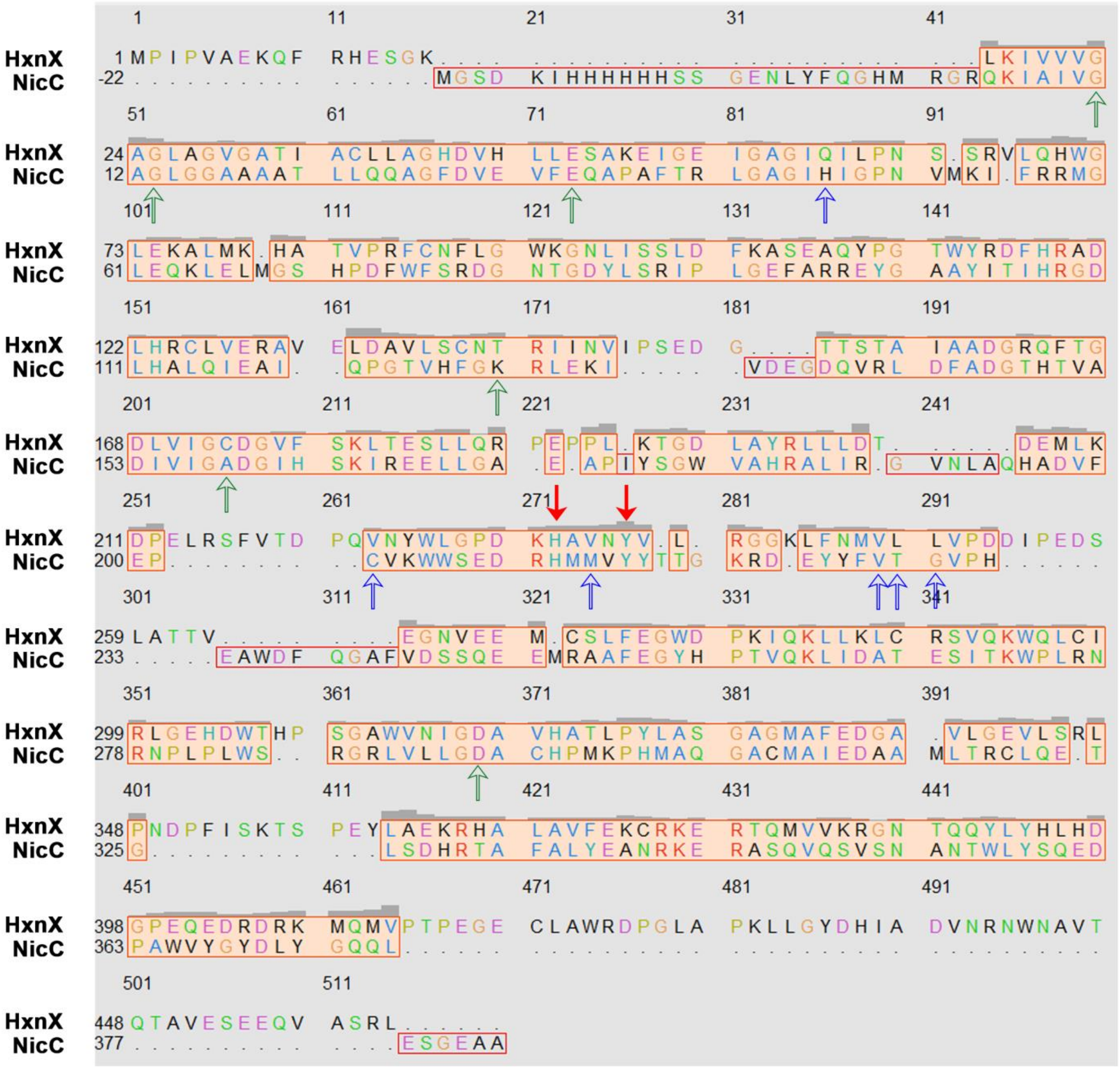

**Fig. S2. HxnX shares structural similarities with the 6-NA 3-monooxygenase, NicC of *P. putida*.** Alignment of the superimposed proteins shown in Fig. 5*A* was conducted by Matchmaker Match Align within Chimera. Solid red arrows indicate the substrate binding residues in NicC; empty red arrows indicate additional AA residues that were mapped to the active centre in NicC; green empty arrows mark the FAD binding residues in NicC (7).

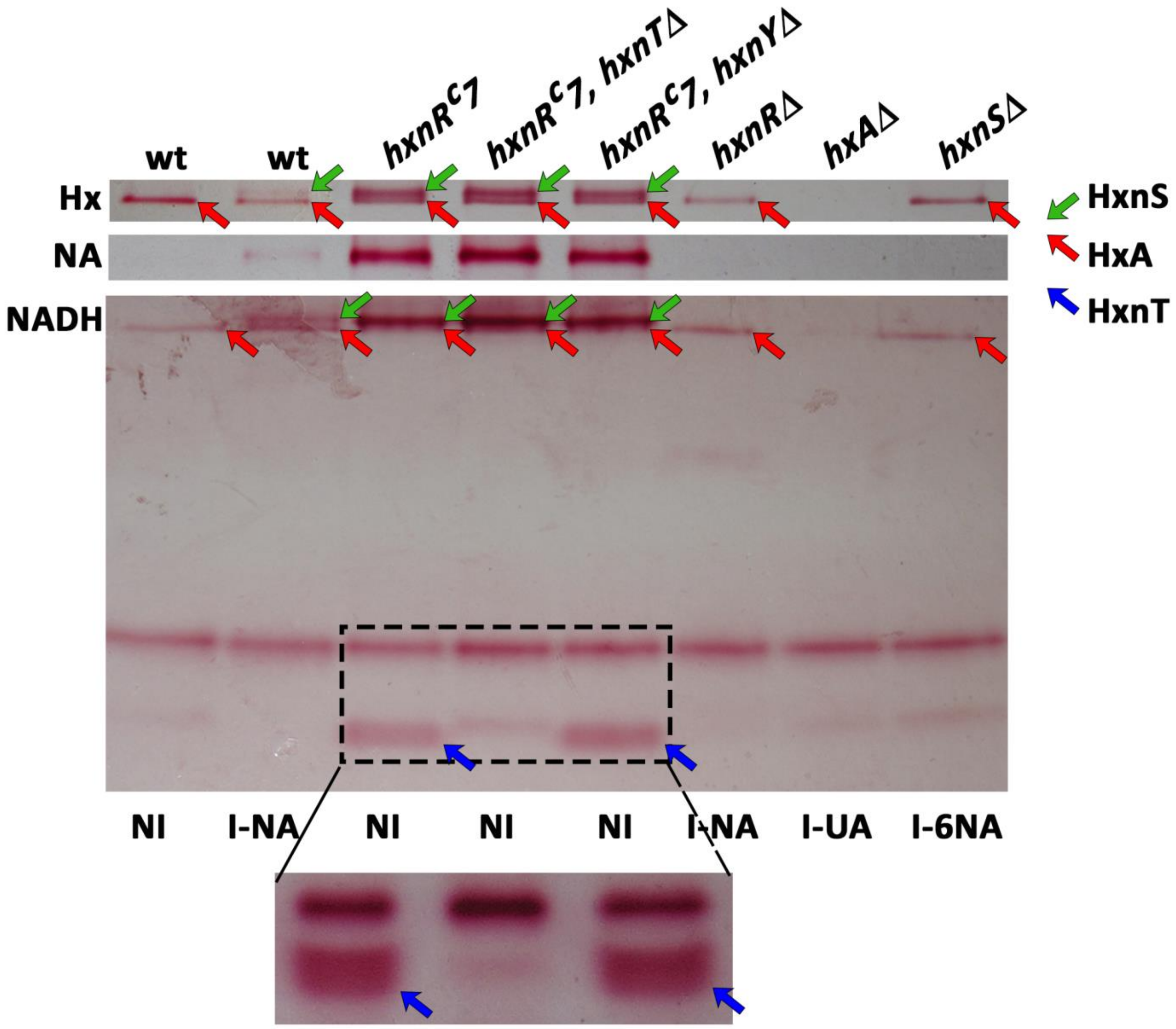


**Fig. S3. Enzyme staining of HxnT with NADH as an electron donor in polyacrylamide gels confirms that HxnT is a FMN flavin-oxidoreductase.**

We have previously shown that both HxA (8) and HxnS (9) can transfer electrons from NADH to tetrazolium. This is shown in the NADH stained native 10% PAGE, where HxnS is shown by an NA inducible band (green arrows) in the wild type (wt) and a strong constitutive band in all *hxnR$^c$7* strains, while a basal level of HxA is shown in all non-induced conditions (without uric acid induction), except in *hxAΔ* strain, from which HxA is missing (red arrows). Two additional, constitutive NADH oxidoreductase bands are seen in all strains. Just below the higher mobility band, a very faint staining band, a new band is seen in *hxnR$^c$7* strains, but not in the *hxnR$^c$7 hxnTΔ* strains (blue arrows). Samples in the boxed area were reloaded in 6-

18 % gradient acrylamide gel (shown below), where the separation of the HxnY specific band from the constitutive band was clearer. NI, non-induced, I-NA, induced with nicotinic acid, I-UA induced with uric acid (which induces HxA but not HxnS), I-6NA, induced with 6-hydroxynicotinic acid. Strains: wt (HZS.145); *hxnR*$^{c}$*7* (FGSCA872); *hxnR*$^{c}$*7 hxnTΔ* (HZS.427); *hxnRΔ* (HZS.614); *hxAΔ* (HZS.245); *hxnSΔ* (HZS.254). Strains were grown on 1 mM acetamide N-source for 20 hours (NI) or 1 mM NA, 1 mM 6-NA or 0.6 mM UA was added to the media at 15 hours for induction. Green, red and blue arrows mark HxnS, HxA and HxnT, respectively.

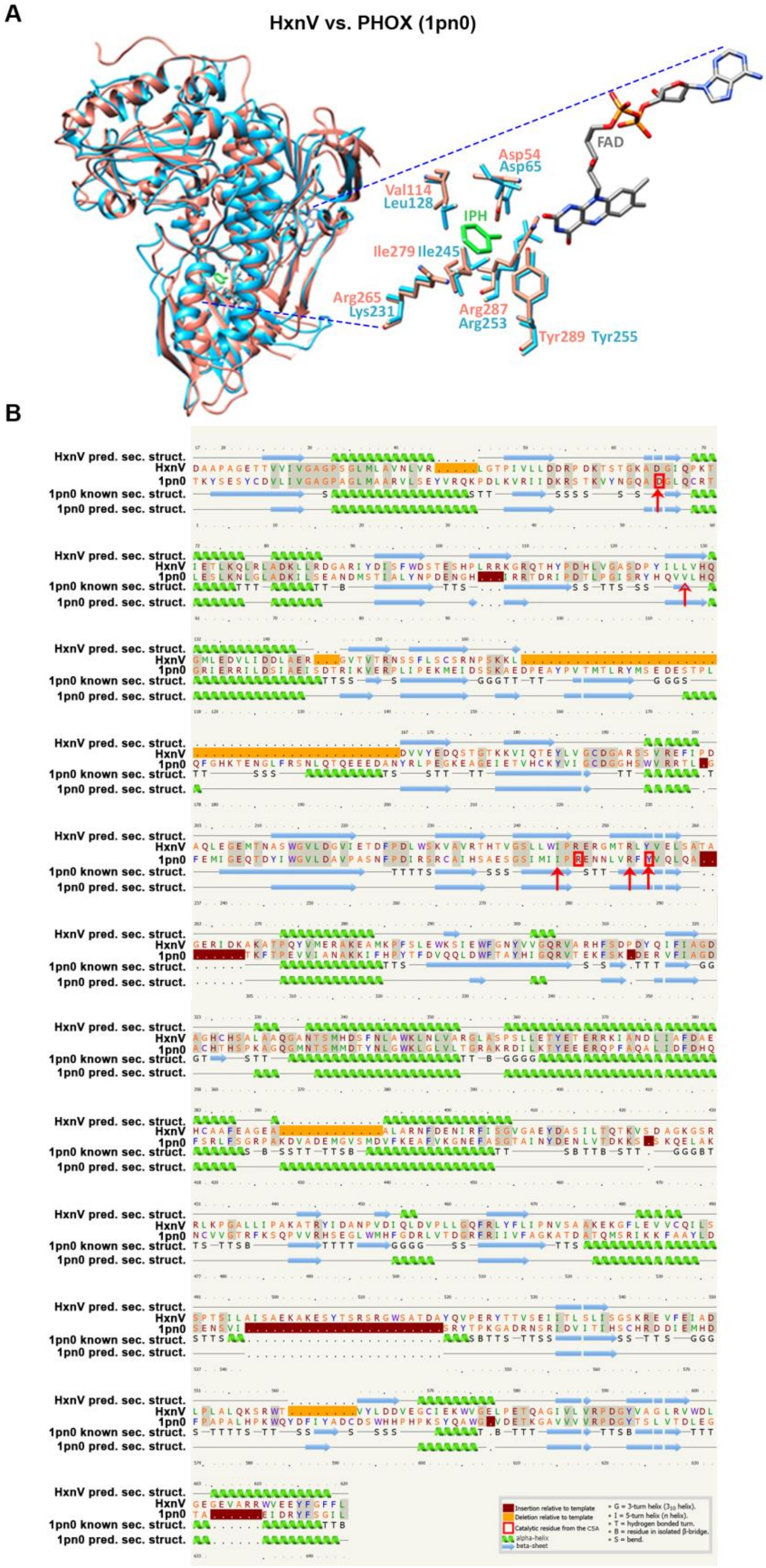


**Fig. S4. Structural comparison of HxnV and the phenol hydroxylase enzyme PHOX from *Trichosporon cutaneum*.** (*A*) Superposition of HxnV with PHOX (PDB code: 1pn0)

shows that functionally important residues are conserved in HxnV. Salmon color shows PHOX, blue color shows HxnV. The Tyr289, Ile279 and Val114 residues of PHOX (Tyr255, Ile245 and Leu128 in HxnV, respectively) establish hydrophobic interaction with the 2- and 6-carbons of the phenol ring, while Tyr289 and Asp54 (Tyr255 and Asp65 in HxnV) form hydrogen bonds with the hydroxyl group of the phenol molecule (IPH, green sticks). Tyr289 (Tyr255 in HxnV) also directly interacts with FAD (10, 11). The Arg287 and Arg265 in PHOX (Arg253 and Lys231 in HxnV) are part of the active site through their interactions with Ile279 and Tyr289 (Ile245 and Tyr255 in HxnV). For quality assessment of the structural model of HxnV, *see Sl Appendix* Table S4. (*B*) The predicted secondary structure of HxnV is shown in comparison both to the known and the predicted secondary structures of PHOX (PDB code: 1pn0) from *T. cutaneum*. Figure was obtained with Phyre2 (1). Red arrows indicate the Phenol-binding AA residues.

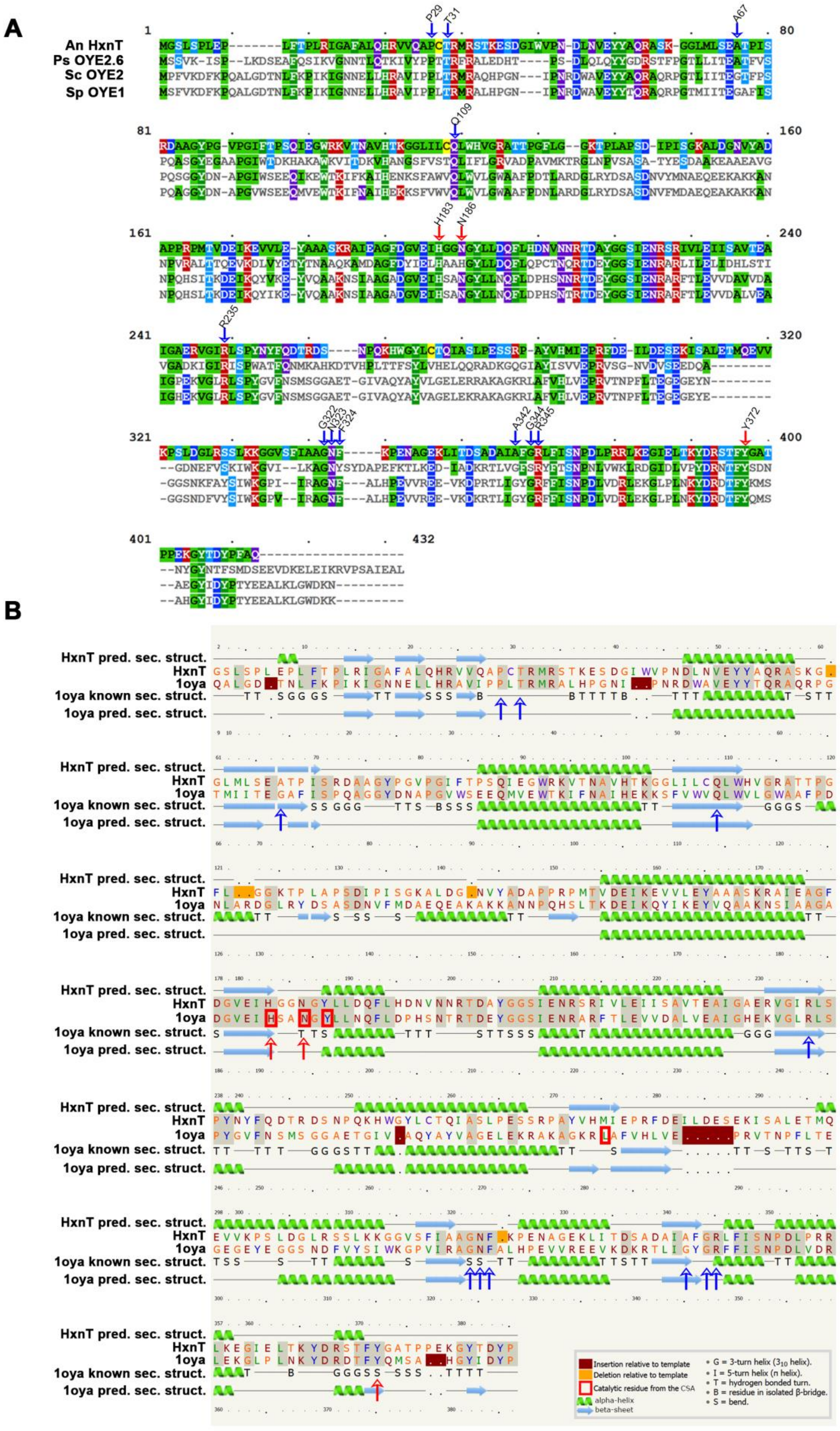

**Fig. S5. Comparison of HxnT to old yellow enzymes.** (*A*) Alignment of the primary structure of HxnT with its nearest homolog enzymes belonging to the OYE-like alkene reductases: PsOYE2.6 from *Pichia stipitis* (XP_001384055; 31.9% identity) (12), ScOYE2 from *Saccharomyces cerevisiae* (AAB68024; 36,8% identity) (13) and SpOYE1 from *Saccharomyces pastorianus* (Q02899; 40.7% identity) (14). Alignment was carried out with Muscle, and visualised with MView. Red and blue arrows indicate equivalent residues of the substrate- and FMN binding sites in HxnT, respectively. FMN binding sites are highly conserved in HxnT. Full conservation of the substrate binding residues of SpOYE1 and ScOYE2 to the corresponding AA residues of HxnT (His/Asn/Tyr AAs numbered as 191/194/375 in SpOYE1, 192/195/376 in ScOYE2 and 183/186/372 in HxnT, respectively) indicates functional similarities of HxnT and ScOYE2 (13) and SpOYE1 (14) rather than with PsOYE2.6 (12). The His/Asn/Tyr triad was shown to establish hydrogen bonds with the nicotinamide moiety of NAD(P)H, which while is quite unusual for most NAD(P)H dependent enzymes is typical of the OYE family (12, 14). *In vitro* NADPH-tetrazolium enzyme assay on *hxnR$^c$7 hxnT$^+$* as control and *hxnR$^c$7 hxnTΔ* strains (see *Sl Appendix* Fig. S8) verified that HxnT can transfer electrons from NADH to tetrazolium, thus HxnT can acts as an FMN flavin-oxidoreductase. (*B*) The predicted secondary structure of HxnT is shown in comparison to both the known and predicted secondary structures of SpOYE1 from *S. pastorianus* (PDB: 1oya). The image was obtained with Phyre2 (1). Red and blue arrows indicate the substrate binding and FMN binding AA residues in SpOYE1 as it is shown in panel *A*.

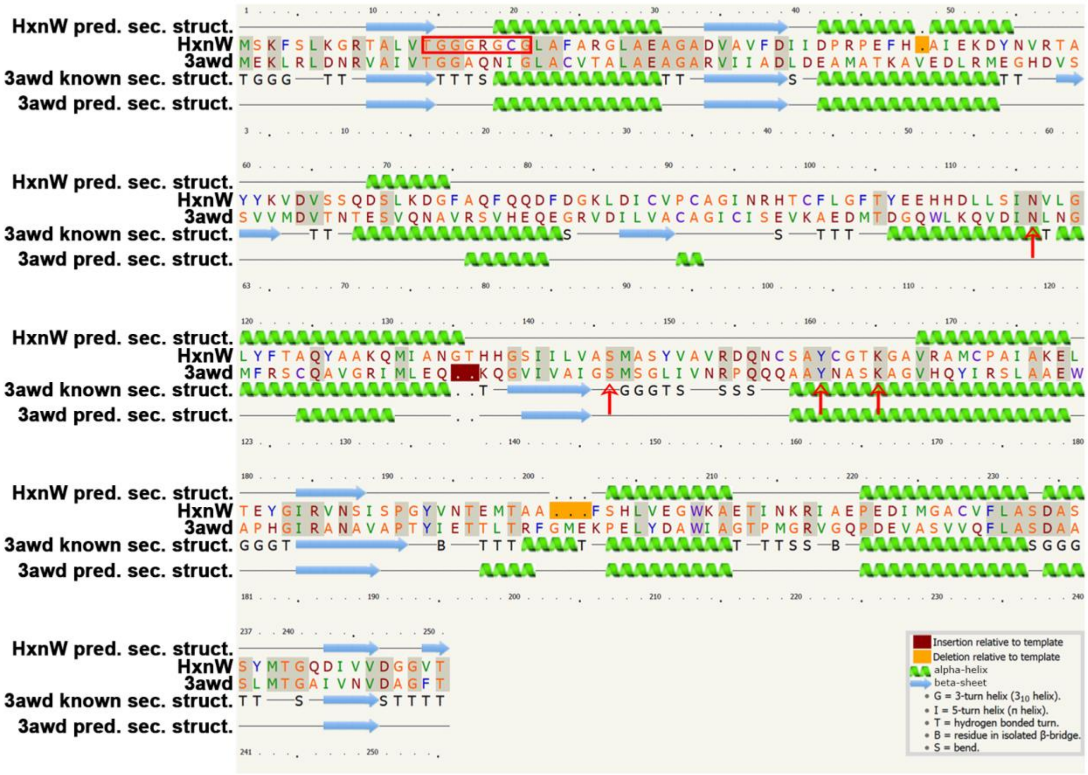

**Fig. S6. Comparison of the predicted secondary structure of HxnW with the secondary structure of Gox2181 from *Gluconobacter oxydans*.** The closest known structural homolog of HxnW is the polyol dehydrogenase enzyme Gox2181 from *G. oxydans* (PDB code: 3awd) (15). The comparative analysis and the image was obtained with Phyre2 (1). Boxed residues show the TG(X)$_3$GXG NAD(P)-binding motif characteristic to the fungal type ketoreductases. Red arrows indicate the conserved catalytic tetrad of the NADB_Rossmann fold domain.

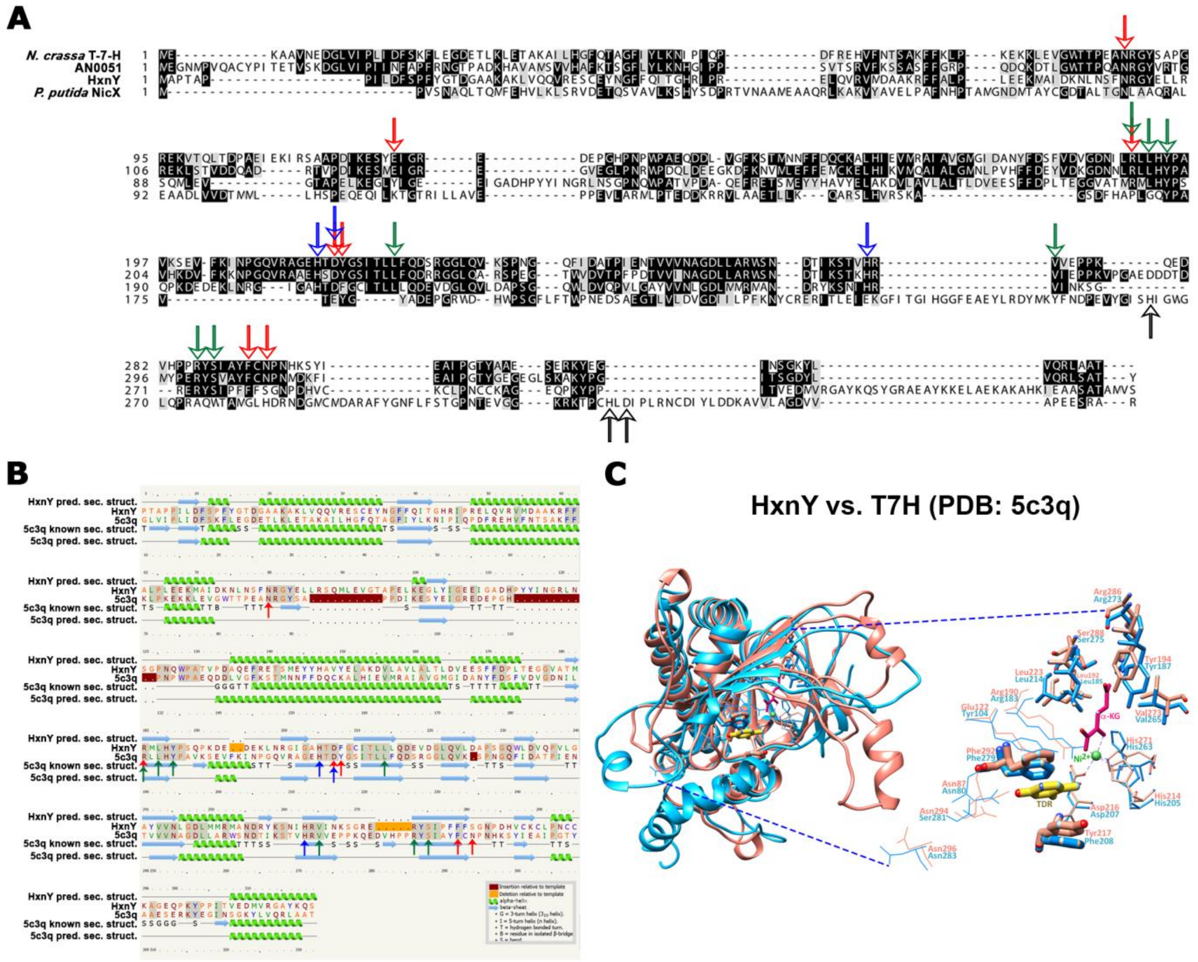


**Fig. S7. Comparative *in silico* analysis of HxnY.** HxnY is an α-ketoglutarate dependent dioxygenase that shares, 28.5% identity with the well-studied thymine-7-hydroxylase (T7H) of *Neurospora crassa* (16), 29.3% with its *A. nidulans* orthologue AN0051, but only 15% with the *A. nidulans* XanA and with the 2,5-dihydroxypyridine dioxygenase of *P. putida* KT2440 (NicX) (17), the latter catalysing the pyridine ring opening in the NA catabolism of this organism. (*A*) Sequence comparison of HxnY with Thymine-7-hydroxylase (T7H) from *N. crassa* (locus NCU06416) (16), its orthologue from *A. nidulans* (AN0051) and NicX, the 2,5-dihydroxypyridine hydroxylase (Q88FY1) from *P. putida* (17). Alignment was carried out with Mafft G-INS-i, and visualised with Box shade. Red, blue and green arrows indicate the thymine binding, α-ketoglutarate binding and Fe(II) binding AA residues from T7H, respectively. Black arrows indicate the Fe(II) binding AA residues in NicX. (*B*) The predicted secondary structure of HxnY in comparison to both the known and the predicted secondary structures of Thymine-7-hydroxylase (T7H) from *N. crassa* (PDB: 5c3q). Image was obtained with Phyre2 (1). Red, blue and green arrows as in Panel *A*. (*C*) Superposition of the structure of HxnY (for quality assessment of the model, see *Sl Appendix* Table S4) with T7H of *N. crassa* (PDB code: 5c3q) (16). Salmon color shows T7H, blue color shows HxnY. TDR: thymine ligand of T7H (yellow thick sticks); $Ni2^{+}$: nickel ion (green sphere); α-KG: α-ketoglutarate ligand (magenta sticks). Thick sticks and medium-sized sticks are residues that establish π-π stacking interactions and direct or indirect hydrogen bonds with TDR ligand in T7H (16). Residues with thin sticks coordinate the metal ion, while wires indicate residues that interact with the α-KG ligand.

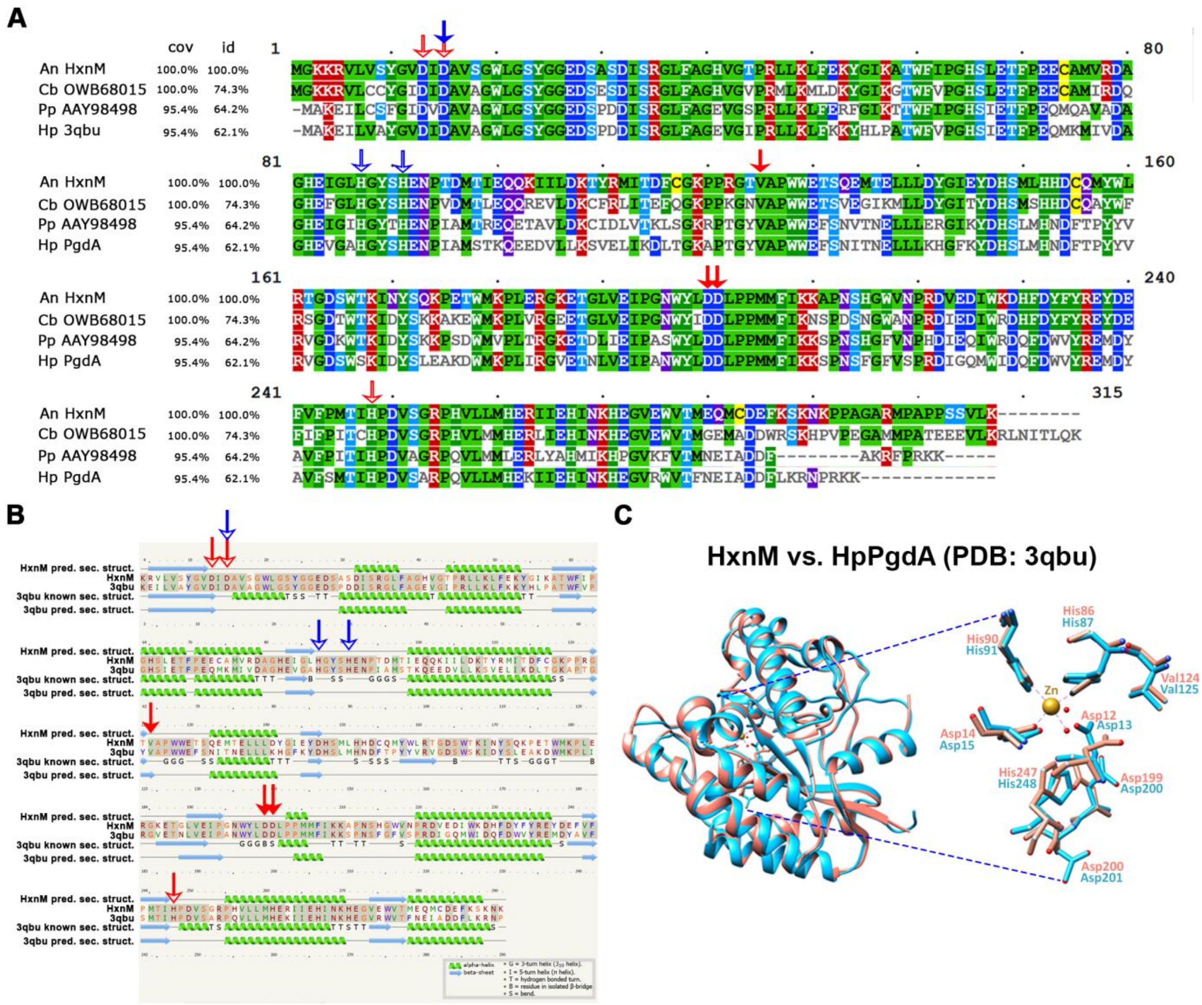


**Fig. S8. Comparison of HxnM to its homologs.** (*A*) Primary sequence comparison of HxnM with the *Candida bondii* hydrolase (OWB68015), the cyclic imide hydrolase of *P. putida* (AAY98498) and HpPgdA of *H. pylori* (18). Level of identity is indicated following protein names as % of identical amino acids and coverage of the aligned sequences. Alignment was carried out with Muscle, and visualised with MView. Hollow red arrows indicate the catalytic water binding AA residues, solid red arrows indicate other catalytic site residues and blue arrows mark the Zn ion coordinating residues in HpPgdA (18, 19). (*B*) HxnM shares structural similarities with the HpPgdA of *H. pylori*. The predicted secondary structure of HxnM is shown in comparison to both the known and the predicted secondary structures of HpPgdA of *H. pylori* (PDB code: 3qbu). Image was obtained with Phyre2 (1) Arrows mark residues as described for panel *A*. (*C*) Superposition of the structural model of HxnM (for quality assessment see *Sl Appendix* Table S4) with its closest known structural homolog, HpPgdA of *H. pylori* (PDI code: 3qbu). Salmon color shows HpPgdA, blue color shows HxnM. The HpPgdA residues His86, His90 and Asp14 (His87, His91 and Asp15 in HxnM, respectively) accomodate the Zn ion (shown in yellow); His247, Asp12 and Asp14 (His248, Asp13 and Asp15 in HxnM, respectively) bind a catalytic water molecule and Asp199, Asp200 and Val124 (Asp200, Asp201 and Val125 in HxnM, respectively) serve other catalytic roles (18). The striking conformity of the active site residues in HxnM indicates functional similarity to HpPgdA.

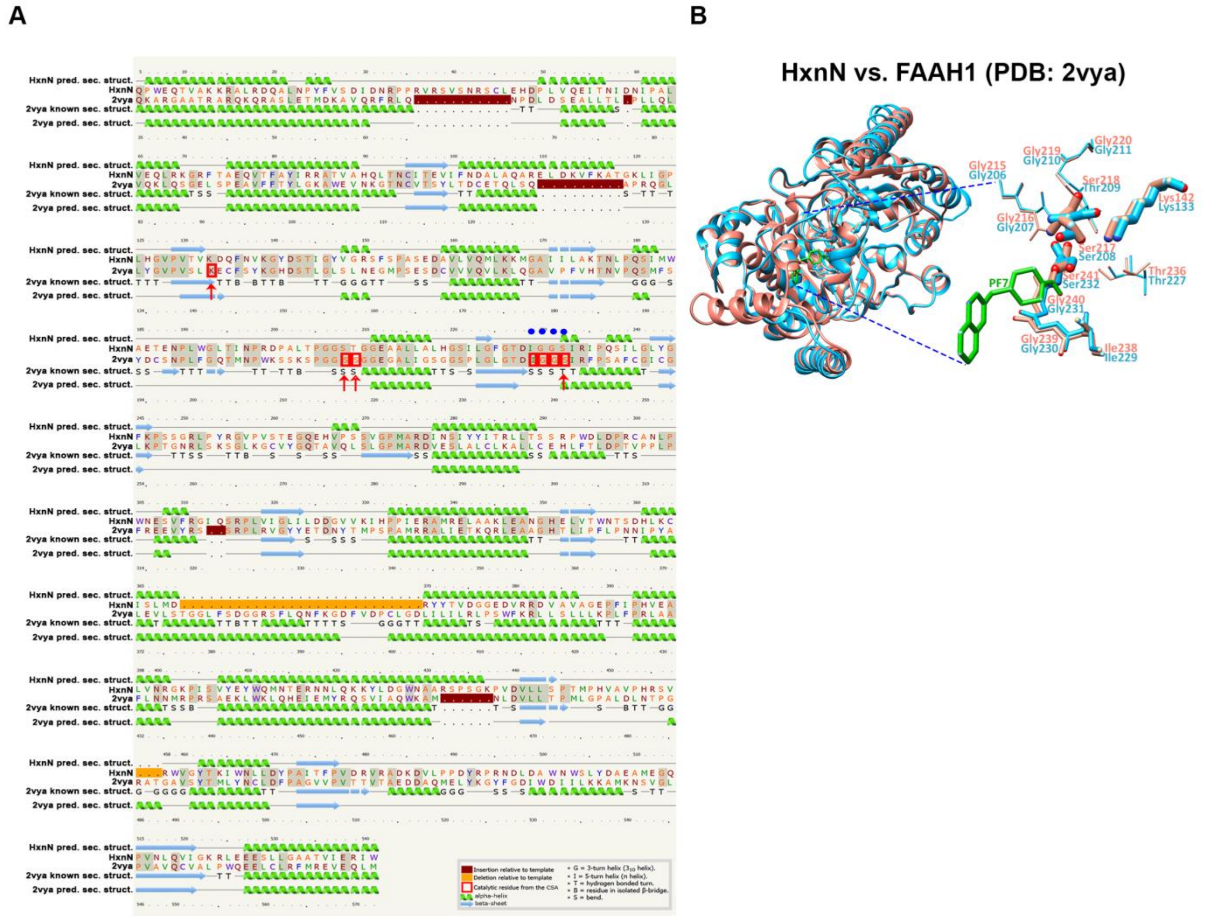


**Fig. S9. HxnN shares structural similarities with the Fatty-acid amide hydrolase 1, FAAH, from *R. norvegicus*.** (*A*) The predicted secondary structure of HxnN is shown in comparison to both the known and the predicted secondary structures of FAAH of *R. norvegicus* (PDB code: 2vya). Image was obtained with Phyre2 (1). Solid red arrows indicate the catalytic triad, the hollow red arrow indicates a Ser that interacts with the catalytic triad and the blue dots indicate the oxyanion hole forming AA residues in FAAH (20, 21). (*B*) Superposition of the structural model of HxnN (for quality assessment see *Sl Appendix* Table S4) with its known structural homolog, fatty acid amide hydrolase 1 (FAAH1) of *Rattus norvegicus* (PDA code: 2vya). Salmon color shows FAAH1, blue color shows HxnN. The Ser217, Ser241 and Lys142 residues in FAAH1 (Ser208, Ser232 and Lys133 in HxnN, respectively) form the catalytic triad for amide bond hydrolysis (thick sticks) (21). The Ile238, Gly239, Gly240 and Ser241 residues in FAAH1 (Ile229, Gly230, Gly231 and Ser232 in

HxnN, respectively) form the oxyanion hole (medium-sized sticks). The catalytic triad in FAAH1 is supported by Gly215, Gly216, Ser218, Gly219, Gly220 and Thr236 residues (thin sticks) (Gly206, Gly207, Thr209, Gly210, Gly211 and Thr227 in HxnN) (20). The 4-(quinolin-3-ylmethyl)piperidine-1-carboxyl acid (PF7) ligand with FAAH1 is shown in green sticks.

HxnN includes a GatA type amidase domain (Asp-tRNAAsn/Glu-tRNAGln amidotransferase A subunit or related amidase domain) (pfam01425) with a transmembrane segment (463-478 AAs) (identified by Phyre2 analysis (1)). Its putative orthologues from *S. cerevisiae* (Amd2p) and *Schizosaccharomyces pombe* (Fah1p) are putative amidases.
