## Supplementary material for "Nicotinate degradation in a microbial eukaryote: a novel, complete pathway extant in *Aspergillus nidulans*": Sl Appendix Table

**Table S1. UHPLC-HRMS characteristics of the intermediates**

| Compound | Elemental compo-sition | RT (min) | Precursor ion | | MS/MS fragments |
|---|---|---|---|---|---|
| | | | Form | Accurate mass | |
| 6-NA* | $C_6H_5NO_3$ | 3.88 | $[M+H]^+$ | 140.0339 | 112.0398, 96.0450, 95.0133, 94.0294, 78.0341, 66.0340 |
| 2,5-DP* | $C_5H_5NO_2$ | 4.92 | $[M+H]^+$ | 112.0400 | 94.0293, 76.0184, 66.0339, 56.0498, 53.0027 |
| 5,6-DHPip-2-O* | $C_5H_9NO_3$ | 4.46 | $[M+H]^+$ | 132.0656 | 115.0397, 114.0556, 97.0293, 87.0447, 86.0607, 71.0494, 69.0337, 59.0495, 55.0183 |
| 3-HPip-2,6-DO* | $C_5H_7NO_3$ | 1.63 | $[M+H]^+$ | 130.0501 | 102.0559, 85.0290, 84.0449, 74.0241, 62.9821, 57.0339 |
| α-HGA* | $C_5H_9NO_4$ | 4.83 | $[M+H]^-$ | 146.0461 | 129.0195, 128.0355, 101.0245, 100.0405, 85.0295, 82.0298, 72.0453 |
| α-HG* | $C_5H_8O_5$ | 3.88 | $[M+H]^-$ | 147.0298 | 129.0194, 103.0401, 101.0244, 85.0295, 57.0341 |

*6-NA: 6-hydroxynicotinic acid; 2,5-DP: 2,5-dihydroxypyridine, 5,6-DHPip-2-O: 5,6-dihydroxypiperidine-2-one, 3-HPip-2,6-DO: 3-hydroxypiperidine-2,6-dione, α-HGA: α-hydroxyglutaramate, α-HG: α-hydroxyglutarate.

**Table S2. NMR results**

**(*A*)**

$^{1}H$, $^{13}C$ (jmod) and 2D NMR spectral assignment for 5,6-dihydroxypiperidine-2-one (500 MHz, $CD_3OD$)
(In $^{1}H$-$^{1}H$ TOCSY spectra all protons have cross-peaks with each other.)

| $\delta_C$(jmod) | | | $\delta_H$ | HMBC | |
|---|---|---|---|---|---|
| 1 | 174.7(+) | | - | | 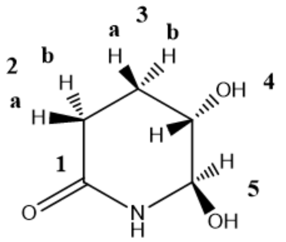 |
| 2 | 27.6(+) | a | 2.26(ddd) | C1, C3, C4 | |
| | | b | 2.50 (ddd) | C1, C3, C4 | |
| 3 | 23.7(+) | a | 2.14(dddd) | C1, C2, C4, C5 | |
| | | b | 1.80(dddd) | C1, C2, C4, C5 | |
| 4 | 68.1(–) | | 3.79(ddd) | C2, C3, C5 | |
| 5 | 80.2(–) | | 4.74 (d) | C1, C3, C4 | |

**(*B*)**

$J_{H\text{-}H}$ couplings ($^{2}J$ and $^{3}J$) for 5,6-dihydroxypiperidine-2-one (500 MHz, $CD_3OH$)

| | 2a | 2b | 3a | 3b | 4 | 5 |
|---|---|---|---|---|---|---|
| 2a | - | *18.0*[a] | 6.5 | 3.5 | | |
| 2b | *18.0*[a] | - | 10.7 | 6.7 | | |
| 3a | 6.5 | 10.7 | - | *13.7*[a] | 2.6 | |
| 3b | 3.5 | 6.7 | *13.7*[a] | - | 5.3 | |
| 4 | | | 2.6 | 5.3 | - | 3.0 |
| 5 | | | | | 3.0 | - |

[a] marks $^{2}J$

**(*C*)**

$^{1}H$, $^{13}C$ (jmod) and 2D NMR spectral assignment for α-hydroxyglutaramate (500 MHz, DMSO)
(Based on $^{1}H$-$^{1}H$ NOESY the amide protons (0a/0b) are closer to 2a/2b than 3a/3b and 4.)

| | $\delta_C$(jmod) | | $\delta_H$ | $^{1}H$-$^{1}H$ COSY | $^{1}H$-$^{1}H$ TOCSY | HMBC | |
|---|---|---|---|---|---|---|---|
| 0 | | a | 6.58(s) | 0b | 0b | | 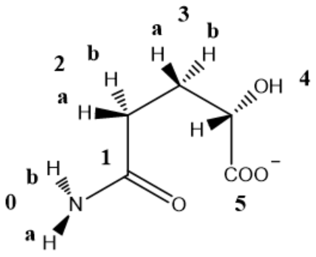 |
| | | b | 7.26(s) | 0a | 0a | | |
| 1 | 174.9(+) | | - | | | | |
| 2 | 31,7(+) | a | 2.04(m) | 2b, 3a, 3b | 2b, 3a, 3b, 4 | C1, C3, C4 | |
| | | b | 2.08(m) | 2a, 3a, 3b | 2a, 3a, 3b, 4 | C1, C3, C4 | |
| 3 | 30.8(+) | a | 1.81(m) | 2a, 2b, 3b, 4 | 2a, 2b, 3b, 4 | C1, C2, C4, C5 | |
| | | b | 1.56(m) | 2a, 2b, 3a, 4 | 2a, 2b, 3a, 4 | C1, C2, C4, C5 | |
| 4 | 70.3(–) | | 3.51(dd)($^{3}J$ = 4.4 / 6.8) | 3a, 3b | 2a, 2b, 3a, 3b | C2, C3, C5 | |
| 5 | 176.4(+) | | - | | | | |

**Table S3. List of *A. nidulans* strains used in this work.** All strains listed are *veA1* mutants.

| Strain | Genotype | Purpose | Reference |
|---|---|---|---|
| A148 | *pabaA1 wA3* | parental strain in genetic crosses with TN02 A21 | provided by Herb Arst |
| CS2638 | *yA2 pantoB100 riboB2 nicB8 fpaD43 acet⁻* | parental strain in genetic crosses with HZS.614, HZS.221 and HZS.226 | provided by S. Amillis |
| FGSCA26 | *biA1* | mRNA expression analysis | (1) |
| FGSCA872/ CS51 | *hxnR$^c$7 biA1* | growth test; enzyme assay; parental strain in genetic crosses with HZS.293, HZS.294 and HZS.288; metabolite analysis | (2) |
| TN02 A21 | *riboB2 pyroA4 nkuAΔ::argB$^+$* | recipient strain in transformation experiment to obtain *hxnP, hxnV, hxnN, hxnR* and *hxnZ* deletions; growth test | (3) |
| NA1322 | *acuL::GFP(at Locus acuL) pabaA1 biA1 argB2; in trans pDsRed-SKL-argB$^+$* plasmid in 1 copy | parental strain in genetic crosses with HZS.305 | (4) |
| HZS.98 | *pantoB100 pabaA1* | parental strain in genetic crosses with HZS.222 | (5) |
| HZS.106 | *hxnSΔ::zeo hxAΔ::zeo pyrG89 pantoB100 biA1 pyr4 in trans* | parental strain in genetic crosses with HZS.122 | (5) |
| HZS.120 | *riboB2 pabaA1* | growth test; recipient strain in transformation experiment to obtain *hxnT* and *hxnY* deletions | (6) |
| HZS.122 | *riboB2 pabaA1 yA2* | parental strain in genetic crosses with HZS.106 | (5) |
| HZS.123 | *anA1 riboB2 pabaA1* | parental strain in genetic crosses with HZS.307 and HZS.393 | this work |
| HZS.143 | *pabaA1 yA2* | parental strain in | this work |

| | | | |
|---|---|---|---|
| | | genetic crosses with HZS.309 | |
| HZS.145 | *veA1* | enzyme assay | (5) |
| HZS.221 | *hxnPΔ::riboB⁺ riboB2 pyroA4 nkuAΔ::argB⁺* | growth test; parental strain in genetic crosses with HZS.399 and CS2638; recipient strain in transformation experiment to obtain *hxnP* and *hxnZ* double deletion mutants | this work (obtained by transformation of the "uphxnP-riboB⁺-downhxnP" substitution cassette into TN02 A21) |
| HZS.222 | *hxnTΔ::pabaA⁺ pabaA1 riboB2* | growth test; parental strain in genetic crosses with HZS.397, HZS.223, HZS.726 | this work (obtained by transformation of the "uphxnT-pabaA⁺-downhxnT" substitution cassette into HZS.120) |
| HZS.223 | *hxnYΔ::riboB⁺ riboB2 pabaA1* | growth test; parental strain in genetic crosses with HZS.395, HZS.222, HZS.548 and HZS.568 | this work (obtained by transformation of the "uphxnY-riboB⁺-downhxnY" substitution cassette into HZS.120) |
| HZS.226 | *hxnZΔ::riboB⁺ riboB2 pyroA4 nkuAΔ::argB⁺* | growth test; parental strain in genetic crosses with HZS.399 and CS2638 | this work (obtained by transformation of the "uphxnZ-riboB⁺-downhxnZ" substitution cassette into TN02 A21) |
| HZS.227 | *riboB2, pantoB100, yA2* | parental strain in genetic crosses with HZS.308 | this work |
| HZS.245 | *hxAΔ::zeo riboB2 pantoB100 biA1* | enzyme assay | (5) |
| HZS.251 | *riboB2 biA1 pabaA1* | recipient strain in transformation experiment to obtain *hxnM* deletion | this work (obtained by genetic cross of HZS.106 with HZS.122) |
| HZS.254 | *hxnSΔ::zeo biA1 pyr4 in trans* | enzyme assay | (5) |
| HZS.267 | *riboB2 pantoB100* | recipient strain for transformation experiment to obtain *hxnM* and *hxnW* deletions | this work (obtained by genetic cross of HZS.308 with HZS.227) |
| HZS.288 | *hxnNΔ::riboB⁺ pyroA4 nkuΔ::argB⁺ riboB2* | growth test; parental strain in genetic crosses with | this work (by transformation of the "uphxnN-riboB⁺- |

| | | | |
|---|---|---|---|
| | | FGSC A872 | downhxnN" substitution cassette into TN02 A21) |
| HZS.292 | *hxnMΔ::riboB*$^+$ *biA1 pabaA1 riboB2* | parental strain in genetic crosses with HZS.294 and HZS.393 | this work (by transformation of the "uphxnM-riboB$^+$-downhxnM" substitution cassette into HZS.251) |
| HZS.293 | *hxnMΔ::riboB*$^+$ *pantoB100 riboB2* | growth test; parental strain in genetic crosses with FGSC A872 and HZS.297 | this work (by transformation of the "uphxnM-riboB$^+$-downhxnM" substitution cassette into HZS.267) |
| HZS. 294 | *hxnVΔ::riboB*$^+$ *pyroA4 nkuΔ::argB*$^+$ *riboB2* | parental strain in genetic crosses with HZS.292 and FGSC A872, growth test | this work (by transformation of the "uphxnV-riboB$^+$-downhxnV" substitution cassette into TN02 A21) |
| HZS.296 | *hxnXΔ::riboB*$^+$ *biA1 pabaA1 riboB2* | parental strain is genetic crosses with HZS.316 | this work (by transformation of the "uphxnX-riboB$^+$-downhxnX" substitution cassette into HZS.251) |
| HZS.297 | *hxnXΔ::riboB*$^+$ *biA1 pabaA1 riboB2* | parental strain in genetic crosses with HZS.293 | this work (by transformation of the "uphxnX-riboB$^+$-downhxnX" substitution cassette into HZS.251) |
| HZS.305 | *hxnXΔ::riboB pantoB100 biA1 (riboB2)* | parental strain in genetic crosses with NA1322 | this work (obtained by genetic cross of HZS.296 with HZS.227) |
| HZS.306 | *hxnNΔ::riboB*$^+$ *pyroA4 hxnR*$^c$*7* (*nkuΔ::argB*$^+$ *riboB2*) | growth test; metabolite analysis | this work (obtained by genetic cross of HZS.288 with FGSC A872) |
| HZS.307 | *hxnR*$^c$*7 pantoB100 biA1* | parental strain in genetic crosses with HZS.123 and HZS.393 | this work (obtained by genetic cross of HZS.293 with FGSC A872) |
| HZS.308 | *hxnMΔ::riboB*$^+$ *pantoB100 hxnR*$^c$*7* (*riboB2*) | growth test; metabolite analysis | this work (obtained by genetic cross of HZS.293 with FGSC A872) |
| HZS.309 | *hxnVΔ::riboB*$^+$ *riboB2 pyroA4 hxnR*$^c$*7* (*nkuΔ::argB*$^+$) | growth test; metabolite analysis; parental strain in genetic crosses with HZS.143 and HZS.429 | this work (obtained by genetic cross of HZS.294 with FGSC A872) |
| HZS.310 | *hxnXΔ::riboB*$^+$ *hxnR*$^c$*7* (*riboB2 nkuΔ::argB*$^+$) | growth test | this work (obtained by genetic cross of |

| | | | |
|---|---|---|---|
| | | | HZS.296 and HZS.316) |
| HZS.316 | *hxnR$^c$7 pyroA4* (*nkuΔ::argB*$^+$ *riboB2*) | parental strain is genetic crosses with HZS.296 | this work (obtained by genetic cross of HZS.288 and FGSC A872) |
| HZS.393 | *hxnWΔ::riboB*$^+$ *riboB2 pantoB100* | growth test; parental strain in genetic crosses with HZS.292 and HZS.123 | this work (by transformation of the "uphxnW-riboB$^+$-downhxnW" substitution cassette into HZS.267 |
| HZS.395 | *hxnR$^c$7 riboB2 biA1* | parental strain in genetic crosses with HZS.223 | this work (obtained by genetic cross of HZS.307 with HZS.123) |
| HZS. 397 | *hxnR$^c$7 pabaA1 anA1* | parental strain in genetic crosses with HZS.222 and HZS.599 | this work (obtained by genetic cross of HZS.307 with HZS.123) |
| HZS.398 | *hxnR$^c$7 riboB2 pabaA1 biA1* | parental strain in genetic crosses with HZS.795 | this work (obtained by genetic cross of HZS.307 with HZS.123) |
| HZS.399 | *hxnR$^c$7 riboB2 pabaA1* | parental strain in genetic crosses with HZS.221 and HZS.226 | this work (obtained by genetic cross of HZS.307 with HZS.123) |
| HZS.404 | *hxnR$^c$7 riboB2 pantoB100* | recipient strain for transformation experiment to obtain *hxnV hxnW* double deletion, *hxnX hxnW* double deletion and *hxnV hxnW hxnX* triple deletion mutants; parental strain in genetic crosses with HZS.726 and HZS.727 | this work (obtained by genetic cross of HZS.307 with HZS.123) |
| HZS.427 | *hxnTΔ::pabaA*$^+$ *hxnR$^c$7 anA1 pabaA1* | enzyme assay; metabolite analysis; parental strain in genetic crosses with HZS.517, HZS.537, HZS.726, HZS.749, HZS.751 and HZS.783 | this work (obtained by genetic cross of HZS.222 with HZS.397) |
| HZS.429 | *hxnYΔ::riboB*$^+$ *hxnR$^c$7 pabaA1 biA1 riboB2* | parental strain in genetic crosses with HZS.309, HZS.517, HZS.726, HZS.727 and HZS.783; | this work (obtained by genetic cross of HZS.223 with HZS.395) |

| | | | |
|---|---|---|---|
| | | metabolite analysis | |
| HZS.480 | *hxnPΔ::riboB*⁺ *hxnZΔ::pyroA*⁺ *riboB2 pyroA4 nkuAΔ::argB*⁺ | growth test | this work (by transformation of the "uphxnZ-pyroA⁺-downhxnZ" substitution cassette into HZS.221) |
| HZS.502 | *hxnTΔ::pabaA*⁺ *hxnYΔ::riboB*⁺ (*riboB2 pabaA1*) | growth test | this work (obtained by genetic cross of HZS.222 with HZS.223) |
| HZS.517 | *hxnWΔ::riboB*⁺ *pantoB100 hxnR*$^c$*7* (*riboB2*) | growth test; metabolite analysis; parental strain in genetic crosses with HZS.427 and HZS.429 | this work (obtained by genetic cross of HZS.393 and HZS.307) |
| HZS.534 | *hxnXΔ::riboB*⁺ *pantoB100 biA1 pabaA1* (*riboB2*) *in trans pDsRed-SKL-argB*⁺ plasmid in 1 copy | parental strain in genetic crosses with HZS.568, HZS.726 and HZS.727 | this work (obtained by genetic cross of HZS.305 and NA1322) |
| HZS.537 | *hxnVΔ::riboB*⁺ *hxnR*$^c$*7 pabaA1 yA2* (*riboB2 nkuΔ::argB*⁺) | parental strain in genetic crosses with HZS.427, HZS.568; HZS.726 and HZS.727 | this work (obtained by genetic cross of HZS.309 with HZS.143) |
| HZS.548 | *hxnSΔ::pabaA*⁺ *pabaA1 anA1 riboB2* | parental strain in genetic crosses with HZS.223 | this work (obtained by genetic cross of HZS.397 and HZS.599) |
| HZS.558 | *hxnSΔ::pabaA*⁺ *hxnYΔ::riboB*⁺ *anA1 pabaA1 riboB2* | growth test | this work (obtained by genetic cross of HZS.223 and HZS.548) |
| HZS.563 | *nkuAΔ::argB*⁺ *pabaA1 riboB2 pyroA4* | recipient strain for transformation experiment to obtain *hxnX* deletion mutants | this work (obtained from genetic cross of TN02 A21 with A148) |
| HZS.564 | *nkuAΔ::argB*⁺ *pabaA1 riboB2 pyroA4* | recipient strain for transformation experiment to obtain *hxnS hxnT* double deletion mutant | this work (obtained from genetic cross of TN02 A21 with A148) |
| HZS.568 | *hxnSTΔ::pabaA*⁺ *pabaA1 pyroA4 riboB2 nkuAΔ::argB*⁺ | parental strain in genetic crosses with HZS.223, HZS.537 and HZS.623; recipient strain for transformation experiment to obtain *hxnSΔ hxnTΔ hxnR*$^c$*7* double | this work (by transformation of "ruphxnS-pabaA⁺-downhxnT" substitution cassette into HZS.564) |

| | | | |
|---|---|---|---|
| | | deletion mutant | |
| HZS.569 | *hxnSTΔ::pabaA*$^+$ *hxnYΔ::riboB*$^+$ *pyroA4 pabaA1 riboB2* (*nkuAΔ::argB*$^+$) | growth test; recipient strain for transformation experiment to obtain *hxnSΔ hxnTΔ hxnYΔ hxnR*$^c$*7* triple deletion mutant | this work (obtained from genetic cross of HZS.568 with HZS.223) |
| HZS.579 | *hxnXΔ::riboB*$^+$ *biA1 pabaA1* (*riboB2*)*; in trans pDsRed-SKL-argB*$^+$ plasmid in 1 copy*; in trans* pAN-HZS-13 plasmid is 7 copy | fluorescent microscopy | this work (by transformation of the pAN-HZS-13 plasmid into HZS.534) |
| HZS.582 | *hxnMΔ::riboB*$^+$ *hxnXΔ::riboB*$^+$ *pantoB100 (riboB2)* | metabolite analysis | this work (obtained from genetic cross of HZS.293 and HZS.297) |
| HZS.584 | *hxnMΔ::riboB*$^+$ *hxnVΔ::riboB*$^+$ *pabaA1 biA1 (riboB2)* | metabolite analysis | this work (obtained from genetic cross of HZS.294 and HZS.292) |
| HZS.588 | *hxnMΔ::riboB*$^+$ *hxnWΔ::riboB*$^+$ *pabaA1 pantoB100 (riboB2)* | metabolite analysis | this work (obtained from genetic cross of HZS.393 with HZS.292) |
| HZS.592 | *hxnTΔ::pabaA*$^+$ *riboB2 pantoB100 (pabaA1)* | parental strain in genetic crosses with HZS.223 | this work (obtained from genetic cross of HZS.98 with HZS.222) |
| HZS.599 | *hxnSΔ::pabaA*$^+$ *pabaA1 riboB2* | growth test; parental strain in genetic crosses with HZS.397; recipient strain for transformation experiment to obtain *hxnS hxnT* double deletion mutant | (5) |
| HZS.614 | *hxnRΔ::AfriboB*$^+$ *riboB2 pyroA4 nkuAΔ::argB*$^+$ | parental strain in genetic crosses with HZS.281 and CS2638; metabolite analysis; mRNA expression analysis | this work (by transformation of the "uphxnR-AfriboB$^+$-downhxnR" substitution cassette into TN02 A21) |
| HZS.623 | *hxnWΔ::riboB*$^+$ *riboB2 pabaA1* | parental strain in genetic crosses with HZS.568 | this work (obtained by genetic cross of HZS.393 and HZS.123) |
| HZS.726 | *hxnXΔ::riboB*$^+$ *riboB2 pabaA1 pyroA4 nkuAΔ::argB*$^+$ | growth test; parental strain in genetic crosses with | this work (by transformation of the "uphxnX-riboB$^+$- |

| | | | |
|---|---|---|---|
| | | HZS.222, HZS.404, HZS.429 and HZS.537 | downhxnX" substitution cassette into HZS.563) |
| HZS.727 | *hxnXΔ::pabaA*$^+$ *pabaA1 riboB2 pyroA4 nkuAΔ::argB*$^+$ | parental strain in genetic crosses with HZS.222, HZS.404, HZS.429 and HZS.537 | this work (by transformation of the "uphxnX-pabaA$^+$-downhxnX" substitution cassette into HZS.563) |
| HZS.747 | *hxnYΔ::riboB*$^+$ *hxnVΔ::riboB*$^+$ *hxnR*$^c$*7 pyroA4* (*riboB2 nkuAΔ::argB*$^+$) | metabolite analysis | this work (obtained by genetic cross of HZS.429 and HZS.309) |
| HZS.748 | *hxnTΔ::pabaA*$^+$ *hxnVΔ::riboB*$^+$ *hxnR*$^c$*7 anA1 pabaA1* (*riboB2 nkuΔ::argB*$^+$) | metabolite analysis | this work (obtained by genetic cross of HZS.427 and HZS.537) |
| HZS.749 | *hxnVWΔ::riboB*$^+$ *riboB2 hxnR*$^c$*7 pantoB100* | metabolite analysis | this work (by transformation of the "uphxnV-riboB$^+$-downhxnW" substitution cassette into HZS.404) |
| HZS.750 | *hxnXWVΔ::riboB*$^+$ *riboB2 hxnR*$^c$*7 pantoB100* | metabolite analysis | this work (by transformation of the "uphxnV-riboB$^+$-downhxnX" substitution cassette into HZS.404) |
| HZS.751 | *hxnXWΔ::riboB*$^+$ *riboB2 hxnR*$^c$*7 pantoB100* | metabolite analysis | this work (by transformation of the "uphxnW-riboB$^+$-downhxnX" substitution cassette into HZS.404) |
| HZS.783 | *hxnXΔ::pabaA*$^+$ *hxnVΔ::riboB*$^+$ *hxnR*$^c$*7 pyroA4 pabaA1 nkuAΔ::argB*$^+$(*riboB2*) | metabolite analysis | this work (obtained by genetic cross of HZS.727 and HZS.537) |
| HZS.795 | *hxnTΔ::pabaA*$^+$ *hxnYΔ::riboB*$^+$ *pantoB100 pabaA1 riboB2* | parental strain in genetic crosses with HZS.398 | this work (obtained by genetic cross of HZS.223 and HZS.592 (derived from subsequent crosses of HZS.98 and HZS.222)) |
| HZS.798 | *hxnXΔ::riboB*$^+$ *hxnTΔ::pabaA*$^+$ *hxnR*$^c$*7 anA1 pabaA1* (*nkuAΔ::argB*$^+$ *riboB2*) | metabolite analysis | this work (obtained by genetic cross of HZS.427 and HZS.726) |
| HZS.810 | *hxnXΔ::pabaA*$^+$ *hxnYΔ::riboB*$^+$ *hxnR*$^c$*7 pyroA4 biA1 riboB2 pabaA1* (*nkuAΔ::argB*$^+$) | metabolite analysis | this work (obtained by genetic cross of HZS.429 and HZS.726) |

| | | | |
|---|---|---|---|
| HZS.812 | *hxnXΔ::pabaA*$^+$ *hxnR*$^c$*7 pabaA1* (*riboB2 nkuAΔ::argB*$^+$) | metabolite analysis | this work (obtained by genetic cross of HZS.537 and HZS.727) |
| HZS.892 | *hxnSΔ::pabaA*$^+$ *pabaA1 hxnTΔ::riboB*$^+$*riboB2* | growth test | this work (by transformation of the "uphxnT-riboB$^+$-downhxnT" substitution cassette into HZS.599) |
| HZS.894 | *hxnWΔ::riboB*$^+$ *hxnTΔ::pabaA*$^+$ *hxnR*$^c$*7* (*riboB2 pabaA1*) | metabolite analysis | this work (obtained by genetic cross of HZS.427 and HZS.517) |
| HZS.898 | *hxnYΔ::riboB*$^+$ *hxnWΔ::riboB*$^+$ *pantoB100 hxnR*$^c$*7* (*riboB2*) | metabolite analysis | this work (obtained by genetic cross of HZS.429 and HZS.517) |
| HZS.899 | *hxnXΔ::pabaA*$^+$ *hxnVΔ::riboB*$^+$ *hxnTΔ::pabaA*$^+$ *pyroA4 hxnR*$^c$*7* (*pabaA1 riboB2*) | metabolite analysis | this work (obtained by genetic cross of HZS.427 and HZS.783) |
| HZS.901 | *hxnXΔ::pabaA*$^+$ *hxnVΔ::riboB*$^+$ *hxnYΔ::riboB*$^+$ *hxnR*$^c$*7* (*pabaA1 riboB2*) | metabolite analysis | this work (obtained by genetic cross of HZS.429 and HZS.783) |
| HZS.902 | *hxnVWΔ::riboB*$^+$ *hxnTΔ::pabaA*$^+$ *pantoB100 hxnR*$^c$*7* (*pabaA1*) | metabolite analysis | this work (obtained by genetic cross of HZS.427 and HZS.749) |
| HZS.903 | *hxnTΔ::pabaA*$^+$ *hxnYΔ::riboB*$^+$ *pantoB100 hxnR*$^c$*7 pabaA1 riboB2* | metabolite analysis | this work (obtained by genetic cross of HZS.398 and HZS.795) |
| HZS.904 | *hxnXWΔ::riboB*$^+$ *hxnTΔ::pabaA*$^+$ *pantoB100 hxnR*$^c$*7* (*pabaA1 riboB2*) | metabolite analysis | this work (obtained by genetic cross of HZS.427 and HZS.751) |
| HZS.911 | *hxnSTΔ::pabaA*$^+$ *pabaA1 riboB2 pyroA4* (*nkuAΔ::argB*$^+$) + *hxnR*$^c$*7 - pyroA*$^+$ in *in trans* pAN-HZS-17 plasmid in 2 copy | metabolite analysis | this work (by transformation of the pAN-HZS-17 plasmid into HZS.568) |
| HZS.912 | *hxnSTΔ::pabaA*$^+$ *pabaA1 hxnYΔ::riboB*$^+$ *riboB2 pyroA4* (*nkuAΔ::argB*$^+$) + *hxnR*$^c$*7 - pyroA*$^+$ in *in trans* pAN-HZS-17 plasmid in 1 copy | metabolite analysis | this work (by transformation of the pAN-HZS-17 plasmid into HZS.569) |

Parenthetic loci indicate alleles that were present in one of the parents of a cross but have not been tested in the progeny.

Explanation of mutant alleles, which are not described in the text: *nkuAΔ* is the deletion of *nkuA* that is essential for non-homologous end joining of DNA in double-strand break repair (3), acet$^-$ is mutation resulting acetate requirement, *fpaD43* is a mutation in *fpaD* resulting p-fluorophenylalanine resistance (7), *veA1* is a mutation in the *veA* gene resulting profuse conidiation regardless of the presence or absence of light (1), *yA2* is mutation in *yA* resulting yellow conidia (8) and *pyr4* is gene for orotidine 5'-phosphate carboxylase in *N. crassa* (9), which complements *pyrG89* allele of *A. nidulans*. Other gene symbols refer to auxotrophies: *argB2*, arginine; *biA1*,

biotin; *pabaA1*, p-aminobenzoic acid; *pantoB100*, pantothenic acid; *pyroA4*, pyridoxine; *pyrG89* uracil or uridine; *riboB2*, riboflavin and *anA1*, thiamine.
*AfriboB*$^{+}$ is *riboB* selection marker gene from *A. fumigatus* used for gene replacement.
*nicB8* is a mutation in a NA biosynthetic pathway gene (*nicB*) resulting NA auxotrophy.
*hxnR*$^{c}$*7* is a mutation in the *hxnR* gene resulting constitutive expression of *hxnR* (and all *hxn* genes) without induction (5).

### Table S4. Results of protein modelling

| | HxnY | HxnT | HxnX | HxnW | HxnV | HxnM | HxnN |
|---|---|---|---|---|---|---|---|
| C-score of i.model[1] | 0.94 | 1.03 | -0.3 | 1.21 | 0.33 | 1.24 | 0.35 |
| estim. TM-score of i.model[2] | 0.84±0.08 | 0.85±0.08 | 0.67±0.12 | 0.88±0.07 | 0.76±0.1 | 0.88±0.07 | 0.76±0.10 |
| estim. RMSD of i.model[3] | 4.6±3.0 Å | 4.6±3.0 Å | 7.8±4.4 Å | 3.4±2.4 Å | 7.1±4.1 Å | 3.7±2.5 Å | 6.7±4.0 Å |
| TM-score of ref.model to i.model[4] | 0.9893 | 0.9885 | 0.9834 | 0.9934 | 0.9943 | 0.9923 | 0.9902 |
| RMSD of ref.model to i.model[5] | 0.748 Å | 0.941 Å | 1.096 Å | 0.488 Å | 0.700 Å | 0.581 Å | 0.856 Å |
| RAMA[6], % of AAs in favoured region | 89.3% (268 AAs) | 83.5% (269 AAs) | 76.6% (305 AAs) | 92.3% (205 AAs) | 79.1% (432 AAs) | 89.1% (230 AAs) | 85.1% (393 AAs) |
| RAMA[6], % of AAs in allowed region | 9.6% (29 AAs) | 15.8% (51 AAs) | 19.6% (78 AAs) | 6.8% (15 AAs) | 18.1% (99 AAs) | 10.1% (3 AAs) | 14.7% (68 AAs) |
| RAMA[6], % of AAs in disallowed region | 1.0% (3 AAs) | 0.6% (2 AAs) | 3.8% (15 AAs) | 0.9% (2 AAs) | 2.7% (15 AAs) | 0.8% (2 AAs) | 0.2% (1 AA) |
| number of non-Pro, non-Gly residues | 300 AAs | 322 AAs | 398 AAs | 222 AAs | 546 AAs | 258 AAs | 462 AAs |
| structural homologues used for superpositioning (PDB number; reference; host organism[7]; enzyme name[8]) | 5c3q (10) *N. crassa*; T7H | 1oya (11) *S. pastorianus*; SpOYE1 | 5eow; (12); *P. putida*; NicC | 3awd (13); *G. oxydans*; Gox2181 | 1pn0 (14); *T. cutaneum* and PHOX; 2dkh (15) *C. testosteroni*; 3HBH | 3qbu (16); *H. pylori*; HpPgdA | 2vyA (17) *R. norvegicus*; FAAH1 |
| RMSD between pruned atom pairs in superposition | 1.056 Å (229 pruned atom pairs) | 0.793 Å (276 pruned atom pairs) | 0.733 Å (282 pruned atom pairs) | 0.760 Å (217 pruned atom pairs) | 0.760 Å (217 pruned atom pairs) | 0.517 Å (289 pruned atom pairs) | 0.704 Å (421 pruned atom pairs) |
| RMSD between all atom pairs in superposition | 4.082 Å (across all 305 pairs) | 5.551 Å (across all 360 pairs) | 1.993 Å (across all 350 pairs) | 1.946 Å (across all 249 pairs) | 1.946 Å (across all 249 pairs) | 0.536 Å (across all 290 pairs) | 2.602 Å (across all 509 pairs) |
| modelled interacting molecules in the PDB model[9] | $Ni^{2+}$; αKG[9]; TDR[9] | FMN[9]; HBA[9] | FAD[9] | - | FAD[9]; IPH[9] (for 1pn0); 3HB[9] (for 2dkh) | $Zn^{2+}$ | PF7[9] |
| proposed enzyme activity | hydroxylation | OYE enzyme (uses NADP(H) for FMN reduction) | conversion of 6-NA to 2,5-DP | none predicted | conversion of 2,5-DP | opening of a saturated pyridine ring between C-N | deamination of the opened saturated pyridine ring |
| possible substrates of enzymes | 6-NA-derivative with closed pyridine ring | 6-NA derivative with closed pyridine ring | 6-NA | 6-NA derived ring | 2,5-DP | saturated pyridine ring | aliphatic NA derived catabolite with amide group |

[1,2,3,4 and 5] C-score of initial model, estimated TM-score of initial model, estimated RMSD of initial model, TM-score of refined model to initial model and RMSD of refined model to initial model, respectively. Initial models were obtained by I-Tasser (PMID:18215316); Refined models were obtained by ModRefiner (PMID:22098752).

[6] RAMA: Ramachandran plot analysis results on non-Proline and non-Glycine regions derived from using Procheck server (https://servicesn.mbi.ucla.edu/PROCHECK/)

[7] Organisms: *N. crassa*: *Neurospora crassa, S. pastorianus*: *Saccharomyces pastorianus*; *P. putida*: *Pseudomonas putida, G. oxydans: Gluconobacter oxydans, T. cutaneum*: *Trichosporon cutaneum*; *C. testosteroni*: *Comamonas testosteroni*; *H. pylori*: *Helicobacter pylori*; *R. norvegicus*: *Rattus norvegicus*.

[8] Proteins: T7H: Thymine-7-hydroxylase, SpOYE1: Old Yellow Enzyme of *S. pastorianus*, NicC: 6-hydroxynicotinic acid 3-monooxygenase; Gox2181: polyol dehydrogenase of C4-C8 dihydroxy alcohols, PHOX: phenol hydroxylase (phenol 2-monooxygenase), 3HBH: 3-hydroxybenzoate hydroxylase; HpPgdA: proposed cyclic imidase of *H. pylori*; FAAH1: fatty acid amide hydrolase 1 of AS (amidase signature) amidases (with Ser-Ser-Lys catalytic triad).

[9] Modelled interacting molecules: $Ni^{2+}$ (nickel ion), αKG (α-ketoglutarate), TDR (thymine), FMN (flavin mononucleotide), HBA: *p*-hydroxybenzaldehyde; FAD (flavin adenine dinucleotide), IPH (phenol), 3HB (3-hydroxybenzoic acid), $Zn^{2+}$ (zinc ion), PF7 (4-(quinolin-3-ylmethyl)piperidine-1-carboxylic acid).

**Table S5.** List of primers used in this study.

| Primer names | Sequence | Primer number |
|---|---|---|
| **Primers used for gene deletions** | | |
| ***hxnP* deletion strain** | | |
| hxnP upst frw | 5'- caccgagctgtagctcacctgcttgatg -3' | **1.** |
| hxnP upst rev | 5'- gtggagatttataaacggttctgtttgg -3' | **2.** |
| hxnP ribo chim frw | 5'- *ccaaacagaaccgtttataaatctccac*cgtacgtagtgtagattcaggcacattgaagcg -3' | **3.** |
| hxnP ribo chim rev | 5'- *gacccagtccctacattctgctgtctctg*ggaaaactgccatgactactaggtggtgctatc -3' | **4.** |
| hxnP downst frw | 5'- cagagacagcagaatgtagggactgggtc -3' | **5.** |
| hxnP downst rev | 5'- cgtaaaccgtctcgtctcgtctgctgacac -3' | **6.** |
| hxnP upst nest frw | 5'- ctcacgcttgtcgagttcgatttccatgatg -3' | **7.** |
| hxnP downst nest rev | 5'- aagcttgatcagcacacagtggaatagctg -3' | **8.** |
| ***hxnY* deletion strain** | | |
| hxnY upst frw | 5'- catatcaaatcagagaggagtctatactg -3' | **9.** |
| hxnY upst rev | 5'- ggatactcaacgattactgctgtttagg -3' | **10.** |
| hxnY ribo chim frw | 5'- *cctaaacagcagtaatcgttgagtatccc*cgtacgtagtgtagattcaggcacattgaagcg -3' | **11.** |
| hxnY ribo chim rev | 5'- *cattatgctagcttacatgacaacaagtac*ggaaaactgccatgactactaggtggtgctatc -3' | **12.** |
| hxnY downst frw | 5'- gtacttgttgtcatgtaagctagcataatg -3' | **13.** |
| hxnY downst rev | 5'- gttgtgtatctcggtgcgaggctctggtac -3' | **14.** |
| hxnY upst nest frw | 5'- gcacgagacacgtcggaatgtatgcaccag -3' | **15.** |
| hxnY downst nest rev | 5'- ctctgacacgactcctagatagcagcatg -3' | **16.** |
| ***hxnZ* deletion strain** | | |
| hxnZ upst frw | 5'- ctactgagcgagagcataatccgtgccgag -3' | **17.** |
| hxnZ upst rev | 5'- cctaataatgataagtgaggccagacgctg -3' | **18.** |
| hxnZ ribo chim frw | 5'- *cagcgtctggcctcacttatcattattag*gcgtacgtagtgtagattcaggcacattgaagcg -3' | **19.** |
| hxnZ ribo chim rev | 5'- *ccacttatcagcacaaactctctgacacg*ggaaaactgccatgactactaggtggtgctatc -3' | **20.** |
| hxnZ downst frw | 5'- cgtgtcagagagtttgtgctgataagtgg -3' | **21.** |
| hxnZ downst rev | 5'- gttccatcgtacagcatgctgactgcatac -3' | **22.** |
| hxnZ upst nest frw | 5'- cattgatcatgagccgctcgatcaacatac -3' | **23.** |
| hxnZ downst nest rev | 5'- agcagcaggttccaatgactcgaagtgac -3' | **24.** |
| hxnZ pyro chim frw | 5'- *cagcgtctggcctcacttatcattattagg*cagttgagcctgagaccaatgaatac -3' | **25.** |
| hxnZ pyro chim rev | 5'- *ccacttatcagcacaaactctctgacacg*gtcagtttagtagctgaagcgttcttattag -3' | **26.** |
| hxnZ upst nest frw2 | 5'- gaatggagaacggagaatggagactg -3' | **27.** |
| ***hxnT* deletion and *hxnS-hxnT* double deletion** | | |
| hxnT upst frw | 5'- ctgtgcagtcattgcgtcatctgcatacac -3' | **28.** |
| hxnT upst rev | 5'- cgactgtctcagtagactacgtcatgagc -3' | **29.** |
| hxnT paba chim frw | 5'- *gctcatgacgtagtctactgagacagtcg*gcacatagctattacacgtatgtttgagac -3' | **30.** |
| hxnT paba chim rev | 5'- *ctatctgtattctgtgtcgtagtattcatg*gtagtttgcttgaatggctaacgaggcattg -3' | **31.** |
| hxnT downst frw | 5'- gaatactacgacacagaatacagatagac -3' | **32.** |
| hxnT downst rev | 5'- catagtcttaaccacgagacgatcagtaac -3' | **33.** |
| hxnT upst nest frw | 5'- cgtcgtcctttcgctttgcctgtttgtatg -3' | **34.** |
| hxnT downst nest rev | 5'- tctgttctactacaggcagcgagttttgtc -3' | **35.** |
| hxnS r up frw | 5'- gtgtactcgttcatcacgccaaag -3' | **36.** |
| hxnS r up rev | 5'- cttgctgttggagacgacttgg -3' | **37.** |

| | | |
|---|---|---|
| hxnS r paba chim frw | 5'- *ccaagtcgtctccaacagcaag*gcacatagctattacacgtatgtttgagac -3' | **38.** |
| hxnS r up nest frw | 5'- cagttgaggcatcttgatgtgag -3' | **39.** |
| hxnS irup frw | 5'- gatgtccaggaatcggcagctatac -3' | **40.** |
| hxnT upst rev2 | 5'- gtgcatacatcacgactgattggagttg -3' | **41.** |
| hxnT ribochim frw | 5'- *caactccaatcagtcgtgatgtatgcac*cgtacgtagtgtagattcaggcacattgaagcg -3' | **42.** |
| hxnT ribochim rev | 5'- *ctccggtcatacttggtcaactcgatac*ggaaaactgccatgactactaggtggtgctatc -3' | **43.** |
| hxnT down frw2 | 5'- gtatcgagttgaccaagtatgaccggag -3' | **44.** |
| paba int frw2 | 5'- gagacatccatcactttaccatcgcccg -3' | **45.** |
| ***hxnR* deletion** | | |
| hxnR upst frw | 5'- cagattcgaacgtacctggtttggtc -3' | **46.** |
| hxnR upst rev | 5'- gtatgtgcaggcgtgtttcttcatc -3' | **47.** |
| hxnR Afuribo chim frw | 5'- *gaagaaacacgcctgcacatac*caggagtctggctcgtttgatc -3' | **48.** |
| hxnR Afuribo chim rev | 5'- *tcagcaagaccttcatag*cttgcgaccgtctgatctagattac -3' | **49.** |
| hxnR downst frw | 5'- atgaaggtcttgctgaatcttggtg -3' | **50.** |
| hxnR downst rev | 5'- ctcttgcactgaccttggtttcac -3' | **51.** |
| hxnR upst nest frw | 5'- gtccggttggcaagacaaagactc -3' | **52.** |
| hxnR downst nest rev | 5'- accaactctgactcgacaatgctcg -3' | **53.** |
| hxnR prom frw | 5'- tttttttgatatccatctggtgcatacattccgacgtg -3' | **54.** |
| hxnR prom rev | 5'- tttttttgatatcgcgtgaaggctttggagcag -3' | **55.** |
| ***hxnX* deletion** | | |
| hxnX upst frw | 5'- cagcgtcaagtctcatatctatactg -3' | **56.** |
| hxnX upst rev | 5'- ggaattaggaaggatctggataccgg -3' | **57.** |
| hxnX ribo chim frw | 5'- *ccggtatccagatccttcctaattcc*cgtacgtagtgtagattcaggcacattgaagcg -3' | **58.** |
| hxnX ribo chim rev | 5'- *gtgtatcatcttctcgctcacatctg*ggaaaactgccatgactactaggtggtgctatc -3' | **59.** |
| hxnX paba chim frw | 5'- *ccggtatccagatccttcctaattcc*gcacatagctattacacgtatgtttgagac -3' | **60.** |
| hxnX paba chim rev | 5'- *gtgtatcatcttctcgctcacatctg*gtagtttgcttgaatggctaacgaggcattg -3' | **61.** |
| hxnX downst frw | 5'- cagatgtgagcgagaagatgatacac -3' | **62.** |
| hxnX downst rev | 5'- gagacttgtactcttatcgttcgc -3' | **63.** |
| hxnX upst nest frw | 5'- caccttctttgtacccgtgctctg -3' | **64.** |
| hxnX downst nest rev | 5'- aagcttgatcagcacacagtggaatagctg -3' | **65.** |
| ***hxnW* deletion** | | |
| hxnW upst frw | 5'- actcctccacccaccgtcttg -3' | **66.** |
| hxnW upst rev | 5'- gtgttcttggcgagcgatgcc -3' | **67.** |
| hxnW ribo chim frw | 5'- *ggcatcgctcgccaagaacacc*gtacgtagtgtagattcaggcacattgaagcg -3' | **68.** |
| hxnW ribo chim rev | 5'- *ctcgcagtggtacagtgctg*ggaaaactgccatgactactaggtggtgctatc -3' | **69.** |
| hxnW downst frw | 5'- cagcactgtaccactgcgag -3' | **70.** |
| hxnW downst rev | 5'- gatcagactagtgctagtggttactaac -3' | **71.** |
| hxnW upst nest frw | 5'- cagcgtcaagtctcatatctatactg -3' | **72.** |
| hxnW downst nest rev | 5'- ccatgaaatggattgtaaacctcaag -3' | **73.** |
| ***hxnV* deletion** | | |
| hxnV upst frw | 5'- acttctcaaactgctcgcgtcc -3' | **74.** |
| hxnV upst rev | 5'- caagacggtgggtggaggagt -3' | **75.** |
| hxnV ribo chim frw | 5'- *actcctccacccaccgtcttg*cgtacgtagtgtagattcaggcacattgaagcg -3' | **76.** |
| hxnV ribo chim rev | 5'- *cacgaccaaggcttcgcaaac*ggaaaactgccatgactactaggtggtgctatc -3' | **77.** |
| hxnV downst frw | 5'- gtttgcgaagccttggtcgtg -3' | **78.** |
| hxnV downst rev | 5'- ctgagacgtcagtgaggacat -3' | **79.** |

| | | |
|---|---|---|
| hxnV upst nest frw | 5'- caagggtgaactgcttcgctagg -3' | **80.** |
| hxnV downst nest rev | 5'- cgaaattgtttctctgcaactggg -3' | **81.** |
| ***hxnM* deletion** | | |
| hxnM upst frw | 5'- ctcattgtagtagcattcattgtcgc -3' | **82.** |
| hxnM upst rev | 5'- cgacaccgtaggatacgagaac -3' | **83.** |
| hxnM ribo chim frw | 5'- *gttctcgtatcctacggtgtcg*cgtacgtagtgtagattcaggcacattgaagcg -3' | **84.** |
| hxnM ribo chim rev | 5'- *gctcttaaactcatcgcacatctg*ggaaaactgccatgactactaggtggtgctatc -3' | **85.** |
| hxnM downst frw | 5'- cagatgtgcgatgagtttaagagc -3' | **86.** |
| hxnM downst rev | 5'- ggcgaaacttgagtacgagtg -3' | **87.** |
| hxnM upst nest frw | 5'- cgaatcccgcaaagcattctg -3' | **88.** |
| hxnM downst nest rev | 5'- gatgtccgggtattcggtgcag -3' | **89.** |
| ***hxnN* deletion** | | |
| hxnN upst frw | 5'- cgtagaggctgttttcatgtcctg -3' | **90.** |
| hxnN upst rev | 5'- cgtttctttgcgactgtctgctc -3' | **91.** |
| hxnN ribo chim frw | 5'- *gagcagacagtcgcaaagaaacg*cgtacgtagtgtagattcaggcacattgaagcg -3' | **92.** |
| hxnN ribo chim rev | 5'- *ctagccgtttcccaattacctgc*ggaaaactgccatgactactaggtggtgctatc -3' | **93.** |
| hxnN downst frw | 5'- gcaggtaattgggaaacggctag -3' | **94.** |
| hxnN downst rev | 5'- gatgcacagattgtgaaacgattg -3' | **95.** |
| hxnN upst nest frw | 5'- cggagaatatgtggttcgagc 3' | **96.** |
| hxnN downst nest rev | 5'- gctcttaaactcatcgcacatctg -3' | **97.** |
| **Gene specific primers** | | |
| hxnR frw | 5'- caagcgagactttccagacctagac -3' | **106.** |
| hxnR rev | 5'- aatgctccgtatcccacatgaaacac -3' | **107.** |
| hxnP frw | 5'- gttgatcgagcggctcatgatcaatgagtc -3' | **108.** |
| hxnP rev | 5'- gcactcagaacaccagcgatattgcccag -3' | **109.** |
| hxnT frw | 5'- ctttgcgctgcagcaccgtgttgtccaag -3' | **110.** |
| hxnT rev | 5'- ctattgagcgaaagggtagtccgtatag -3' | **111.** |
| hxnY frw | 5'- caagcttgtacagcaagtacgcgaatcctg -3' | **112.** |
| hxnY rev | 5'- cagtttggcagacacttgcacacatggtctg -3' | **113.** |
| hxnZ frw | 5'- cagatcatcatggaaatcgaactcgacaag -3' | **114.** |
| hxnZ rev | 5'- ctgactggatgcagtagttgcggtaggtc -3' | **115.** |
| hxnX frw | 5'- cgacagctattgctgcggac -3' | **116.** |
| hxnX rev | 5'- ctgagacgtcagtgaggacat -3' | **117.** |
| hxnW frw | 5'- gcagatcgtggtgttcttcgtag -3' | **118.** |
| hxnW rev | 5'- ctcgcagtggtacagtgctg -3' | **119.** |
| hxnV frw | 5'- cagcgtcaagtctcatatctatactg -3' | **120.** |
| hxnV rev | 5'- cagagcacgggtacaaagaaggtg -3' | **121.** |
| hxnM frw | 5'- caagacctaccgcatgattacagac -3' | **122.** |
| hxnM rev | 5'- gtttattgatgtgttcgatgatcctaac -3' | **123.** |
| hxnN frw | 5'- cgaatcccgcaaagcattctg -3' | **124.** |
| hxnN rev | 5'- ctctgtccacagggaaagtgatggc -3' | **125.** |
| hxnT prom frw2 | 5'- gattctcggtcatgtagagcagg -3' | **126.** |
| hxnT prom rev2 | 5'- gcctgcatggacaagctatttag -3' | **127.** |
| hxnV AS frw | 5'- cagcgtcaagtctcatatctatactg -3' | **128.** |
| hxnV AS rev | 5'- cagagcacgggtacaaagaaggtg -3' | **129.** |
| hxnW AS frw | 5'- gcagatcgtggtgttcttcgtag -3' | **130.** |
| hxnW AS rev | 5'- ctcgcagtggtacagtgctg -3' | **131.** |
| **For cloning** | | |
| pyroA HindIII frw | 5'-tttttttaagcttcagttgagcctgagaccaatgaatac -3' | **132.** |
| pyroA HindIII rev | 5'-tttttttaagcttgtcagtttagtagctgaagcgttcttattag -3' | **133.** |
| AN11197 prom NotI frw | 5'-tttttttgcggccgccatctggtgcatacattccgacgtg -3' | **134.** |

| | | |
|---|---|---|
| AN11197 term NheI rev | 5'-tttttttgctagccgccggtgagaaacttataacagac -3' | **135.** |
| pgpd int2 frw | 5'- catgaatctgaggactgcaatcgc -3’ | **136.** |
| trpC term rev | 5'- cgatcttatatccagattcgtcaagctg -3’ | **137.** |
| 1pGpd int frw | 5'- cagtatattcatcttcccatccaagaac -3’ | **138.** |
| 10GFP linker hmgB rev | 5'- *atcaagatcgactgtatcaataag*cttgtacagctcgtccatgccgtg -3’ | **139.** |
| linker kim hxnX frw | 5'- *acaagcttattgatacagtcgatcttgat*atgcccatcccagttgcagagaaac -3’ | **140.** |
| 5GFP NcoI start frw | 5'- tttttttccatggtgagcaagggcgaggagc -3’ | **141.** |
| hxnX NotI rev | 5'- tttttttgcggccgctcataaccgcgatgctacctgttc -3’ | **142.** |
| **Sequencing primers** | | |
| AN11197 2F | 5'- gcacgcttatcgtctccactg -3' | **148.** |
| AN11197 800F | 5'- gtatgatgccaatacagtaaagctacc -3' | **149.** |
| AN11197 1375F | 5'- cttctcccgcttcaatactacatacc -3' | **150.** |
| AN11197 1958 F | 5'- agagatacagaacatgcatttcttccc -3' | **151.** |
| **RT-qPCR** | | |
| actin ReTi frw | 5'- ggtatcatgatcggtatggg -3' | **152.** |
| actin ReTi rev | 5'- tatctgagtgtgaggatacca -3' | **153.** |
| hxnR ReTi frw | 5'- cggcttctgttctactacagg -3' | **154.** |
| hxnR ReTi rev | 5'- cagtctaggtctggaaagtctc -3' | **155.** |
| hxnX ReTi frw | 5'- cttgtatcatctccacgacgg -3' | **156.** |
| hxnX ReTi rev | 5'- ggctaaacactctccctctg -3' | **157.** |
| hxnS Reti frw | 5'- gagcatttctatcttgagacga -3' | **158.** |
| hxnS Reti rev | 5'- ccattgtgttctgggtactg -3' | **159.** |
| AN5650 ReTi frw | 5'- atgtctgctgttatctatctgctc -3' | **160.** |
| AN5650 ReTi rev | 5'- gccaatcctccttaccttcc -3' | **161.** |
| actin ReTi frw2 | 5'- accatgtaccctggtatctc -3' | **162.** |
| actin ReTi rev2 | 5'- ggaggagcaatgatcttgac -3' | **163.** |

Underlined letters in the primer sequences or in the primer names refer to the restriction sites designed within.

Italic letters at the 5' end refer to the chimeric nature of the primer.
